# The Typhoid Toxin Produced by the Nontyphoidal *Salmonella* Serovar Javiana Can Utilize Multiple Binding Subunits, which Compete for Inclusion in the Holotoxin

**DOI:** 10.1101/666016

**Authors:** A Gaballa, AS Harrand, AR Cohn, M Wiedmann, RA Cheng

## Abstract

*Salmonella enterica* encodes a wide array of virulence factors. One novel virulence factor, a DNA-damaging toxin known as the typhoid toxin (TT), was recently characterized in >40 nontyphoidal *Salmonella* (NTS) serovars. Interestingly, these NTS serovars, including *S. enterica* subsp. *enterica* serovar Javiana, also encode *artB*, a homolog of the binding subunit (PltB) of the TT. Here, we show that ArtB and PltB compete for inclusion in the pentameric binding subunit of the TT. Using a combination of *in silico* modeling, a bacterial two-hybrid system expressed in *S.* Javiana, and tandem affinity purification (TAP) of the holotoxin subunits, we show that ArtB and PltB interact *in vivo*. Furthermore, binding subunits composed of homo- and heteropentamers of ArtB and PltB are able to associate with CdtB and PltA to form biologically active toxins. As *artB* was, (i) conserved among *S.* Javiana isolates, and (ii) co-expressed with *pltB* and *cdtB* under Mg^2+^-limiting conditions, we hypothesized that ArtB and PltB compete for inclusion in the binding subunit. Using a novel competition assay, we show that PltB outcompetes ArtB for inclusion in the binding subunit, when cultured at neutral pH. Together, our results suggest that the TT produced by *S.* Javiana utilizes multiple configurations of the binding subunit, representing a novel toxin form and adaptation mechanism for the AB_5_ toxin family. Our work suggests that *Salmonella* serovars, including *S.* Javiana, evolved to encode and maintain multiple binding subunits that can be used to form an active toxin, which may enhance the variety of cells, tissues, or hosts susceptible to this novel form of the TT.

## Introduction

The TT produced by *S.* Typhi, and >40 NTS serovars (den Bakker et al., 2011;Miller and Wiedmann, 2016a), has been proposed as a key virulence factor, acting at both the single-cell and systemic levels (Haghjoo and Galán, 2004;Spanò et al., 2008;Song et al., 2013;Del Bel Belluz et al., 2016;Miller et al., 2018). The TT incorporates the nuclease activity of the CdtB subunit of the cytolethal distending toxin (CDT) (Haghjoo and Galán, 2004;Nešić et al., 2004) with the mono-ADP-ribosyltransferase activity of the pertussis toxin (called the PltA subunit), resulting in a toxin that induces a DNA damage response (DDR) and subsequent cell cycle arrest (Spanò et al., 2008;Song et al., 2013) in eukaryotic cells. The resulting damaged DNA is proposed to play a role in both disease manifestation (Song et al., 2013) and colonization and persistence of the toxin-producing bacteria in the host (Ge et al., 2005; Del Bel Belluz et al., 2016). The binding subunit of the TT was originally characterized as a pentameric ring of PltB monomers (Song et al., 2013).

A homolog of PltB, called ArtB, was originally identified in *S.* Typhimurium DT104 (Saitoh et al., 2005). As a proof of concept, Gao et al., showed that ArtB from *S.* Typhimurium DT104 can assemble into a homopentamer *in vitro*, and further confirmed that the ArtB homopentamer can interact with CdtB and PltA from *S.* Typhi and form a biologically active holotoxin both *in vitro* and *in vivo* (Gao et al., 2017). While both ArtB and PltB bind to glycan modifications on sialic acids on host cells, ArtB is able to bind both Neu5Ac- and Neu5Gc-terminated glycans on sialic acids, whereas PltB preferentially binds Neu5Ac-terminated glycans (Song et al., 2013;Gao et al., 2017). Gao et al. proposed that the expanded binding repertoire of ArtB could reflect the expanded host range of *S.* Typhimurium DT104, despite the fact that serovar Typhimurium isolates do not encode *cdtB, pltA*, or *pltB* (Gao et al., 2017).

We previously established that multiple NTS serovars naturally encode both *artB* and *pltB* (den Bakker et al., 2011;Rodriguez-Rivera et al., 2015;Miller et al., 2018), including *S.* Javiana, the 4^th^ most commonly isolated *S. enterica* serovar associated with human clinical illness in the US (CDC, 2016). As deletion of both *artB* and *pltB* is necessary to abolish toxin activity (Miller et al., 2018), we hypothesized that *S.* Javiana isolates have the ability to produce holotoxins with binding subunits composed of homo-or heteropentamers of either ArtB or PltB, or a mixture of both. Here, we show that *S.* Javiana can use both ArtB and PltB as homo- and heteropentameric binding subunits to form a biologically-active toxin, and that both *artB* and *pltB* are co-expressed and compete for inclusion in the binding subunit of the holotoxin.

## Materials and Methods

### Bacterial strains, plasmids, primers, and media

Bacterial strains used in this study are listed in Table S1, vectors and recombinant constructs in Table S2, and primers in Table S3. Bacterial strains were routinely grown in Difco Lennox broth pH 7 (LB; Becton Dickinson [BD], Franklin Lakes, NJ). For two-hybrid system interactions, *E. coli* and *S.* Javiana strains were grown in M63 medium with maltose as the sole carbon source (Battesti and Bouveret, 2012). N-salts minimal medium (pH 5.8 or 7) containing 8 µM MgSO4 (Deiwick et al., 1999) was used for experiments assessing RNA transcript levels. Unless otherwise indicated, ampicillin and kanamycin were used at 100 µg/mL and 50 µg/mL, respectively, in complex medium (LB) and 50 µg/mL and 25 µg/mL, respectively, in chemically defined medium (M9 or N-salts minimal media).

Human intestinal epithelial cells (HIEC-6 cells; ATCC, Manassas, VA), (Perreault and Beaulieu, 1996), were routinely cultured in Opti-MEM (Gibco-Invitrogen, Carlsbad, CA) medium supplemented with 10% (v/v) fetal bovine serum (FBS; Gibco-Invitrogen) and recombinant epidermal growth factor (10 ng/ml; Gibco-Invitrogen) at 37°C with 5% CO_2_. Cell supernatants were tested routinely for *Mycoplasma* spp. and *Acheoplasma* spp. infection using the VenorGEM *Mycoplasma* detection kit (Sigma-Aldrich, St. Louis, MO).

### Cloning and expression of toxin proteins in *E. coli*

To express CdtB, PltA, PltB, and ArtB, we cloned and expressed the following constructs in *E. coli* BTH101 cells (Karimova et al., 2001): (i) *pltB*-*3xFLAG*-*artB*-*c-Myc* cloned into the high copy number pUT18 vector, or (ii) *cdtB*-*His*-*pltA*-*FLAG* and *cdtB-His*-*pltA-Strep*, each cloned into the low copy number pKNT25 plasmid. All PCRs were performed using the high-fidelity polymerase Q5 (New England Biolabs [NEB], Ipswich, MA) and the primers listed in Table S3. All constructs were designed to contain identical ribosome binding sites (AGGAGG) 5-7 bases upstream of the ATG start codons. The DNA sequences of all constructs were confirmed with Sanger sequencing.

Construction of pUT18::*pltB*-*3×FLAG*-*artB*-*c-Myc* was done as follows: PCR products *pltB-3×FLAG* and *artB*-*c-Myc*, were digested using XbaI-KpnI and KpnI-EcoRI, respectively and ligated into XbaI-EcoRI-digested pUT18 using T4 DNA Ligase (NEB) to form pUT18::*pltB*-*3xFLAG*-*artB*-*c-Myc*. For construction of pKNT25::*cdtB*-*His*-*pltA*-*FLAG* and pKNT25::*cdtB-His*-*pltA-Strep* PCR products *cdtB*-*His* and *pltA-FLAG* or *pltA*-*Strep* were digested with XbaI-KpnI and KpnI-SacI respectively, followed by ligation into XbaI-SacI-digested pKNT25. In order to construct pKNT25 containing *cdtB-His* only, the *pltA-FLAG* region was deleted from pKNT25::*cdtB*-*His*-*pltA*-*FLAG* by digesting plasmid DNA with KpnI and SacI. The resulting 3’ overhang was removed, and the 5’ overhang was filled in with T4 DNA polymerase (NEB). Following purification, the plasmid was self-ligated with T4 DNA ligase. pKNT25::*pltA-FLAG*, pUT18::*pltB-3×FLAG* and pUT18::*artB-c-Myc* were constructed similarly.

Expression of these constructs was carried out in an *E. coli* BTH101 cAMP-null strain (*cyaA*^-^) due to the observed toxicity of TT components in *E. coli* NEB5α cells (NEB).Sequence-confirmed clones were co-transformed (pUT18 and pKNT25) into *E. coli* BTH101 cells, and were selected for ampicillin and kanamycin resistance on LB agar + 1% glucose, incubated at 37°C.

### Assessment of two-hybrid system interactions

Interactions of CdtB, PltA, PltB, and ArtB subunits were first assessed using the Bacterial Adenylate Cyclase Two-Hybrid (BACTH) system in *E. coli* BTH101 cells (Karimova et al., 2001). The full-length polypeptides of the toxin subunits (including signal peptides) were fused to the N-terminal of T25 or T18 subunits in pKNT25 and pUT18, respectively (Fig. 3B). To ensure cytoplasmic CyaA activity, we also cloned the coding regions of CdtB, PltA, PltB, and ArtB lacking the signal peptide, into BACTH plasmids for fusion at either the N-terminal (pKNT25, pUT18) or C-terminal (pKT25, pUT18C). Regions encoding the C-terminal interacting domains of PltA, PltB, and ArtB, as predicted from the TT structure (Song et al., 2013;Gao et al., 2017), were also cloned into pKNT25, pUT18, pKT25, and pUT18C plasmids (Table S2 and S3). As 324 theoretical combinations exist (see Table S4), we developed a high-throughput screening method. Screening was performed by co-transforming T18 and T25 vectors (see Table S5 for details) into *E. coli* BTH101 cells, followed by selection on LB agar + 1% glucose with ampicillin and kanamycin, and subsequently sub-streaking onto LB agar with ampicillin and kanamycin plates containing 1 mM IPTG and 40 µg/mL of X-gal for blue white screening. To ensure that positive interactions were not false positive clones arising from cAMP-independent CAP* spontaneous mutations, plasmid DNA was purified from blue colonies, which was retransformed into *E. coli* BTH101 competent cells. Plasmids were isolated from clones that were blue after the second transformation, and were analyzed by Sanger sequencing to determine the identity of the interacting domains.

**Figure 1.**
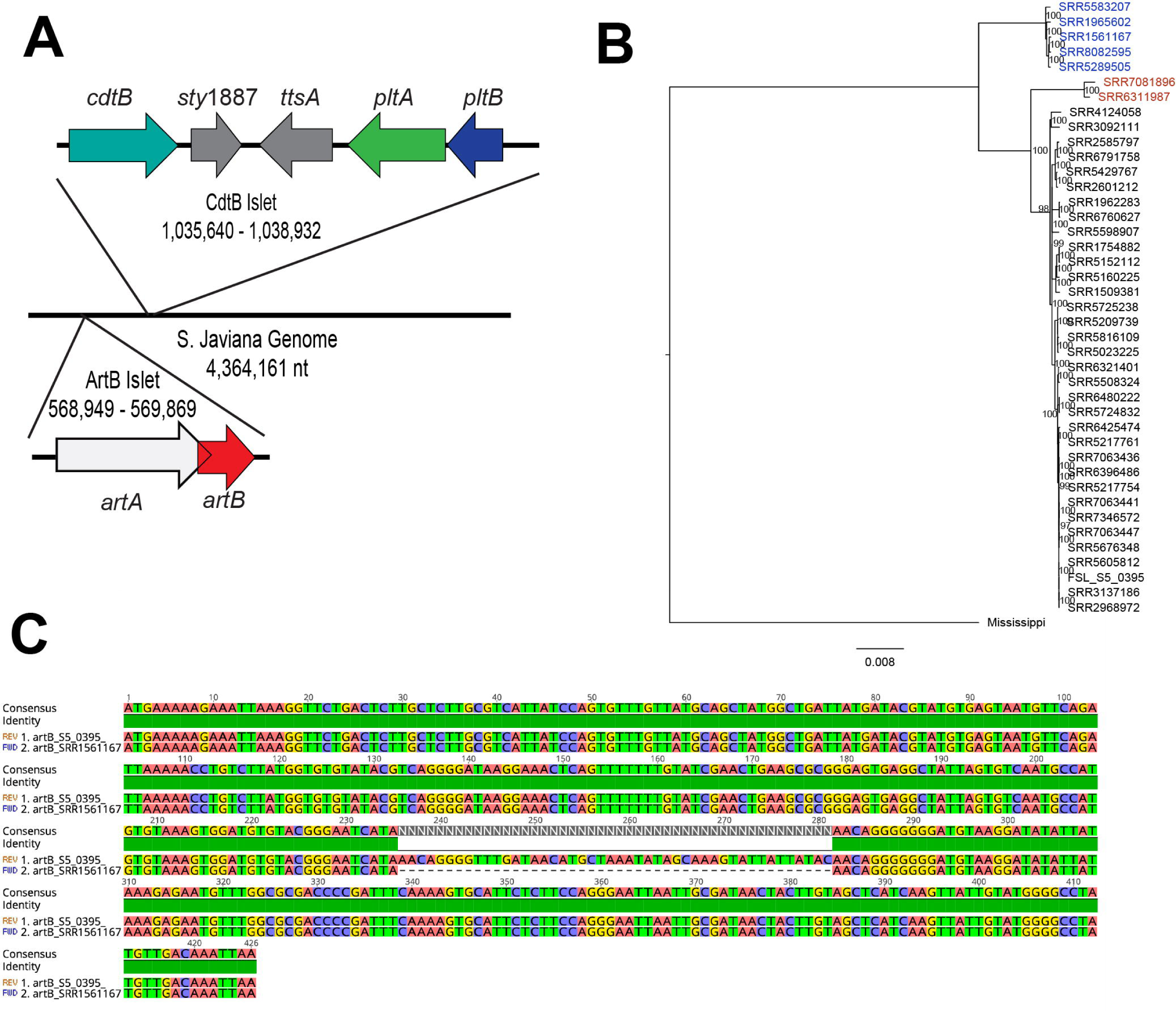
Operon structure and conservation of *artB* in *S.* Javiana. (**A**) Operon structure of the *cdtB*-islet and *artB*-islet in *S.* Javiana strain CFSAN001992. *artA* in *S.* Javiana is a pseudogene as it has a frame shift mutation leading to a premature stop codon; the end of *artA* is predicted to be within *artB* as shown. (**B**) Maximum likelihood tree constructed with core SNPs from *S.* Javiana isolates. RaXML was used to generate the maximum likelihood tree using a general time-reversible model with gamma-distributed sites. One thousand bootstrap repetitions were performed; only bootstrap values >70 are shown. *S.* Mississippi isolate SRR1960042 was used as an outgroup to root the tree. The *S.* Javiana strain (FSL S5-0395) that was used for all experiments is also included in the phylogenetic tree. Isolates for which *artB* was not detected are shown in red and isolates with the 46-nucleotide deletion in *artB* are shown in blue. The scale bar represents the average number of nucleotide substitutions per site. (**C**) DNA sequence alignment between *S.* Javiana strain FSL S5-0395 and a representative isolate (SRR1561167) containing the 46-nucleotide deletion, which results in a frameshift that generates a premature stop codon.

**Figure 2.**
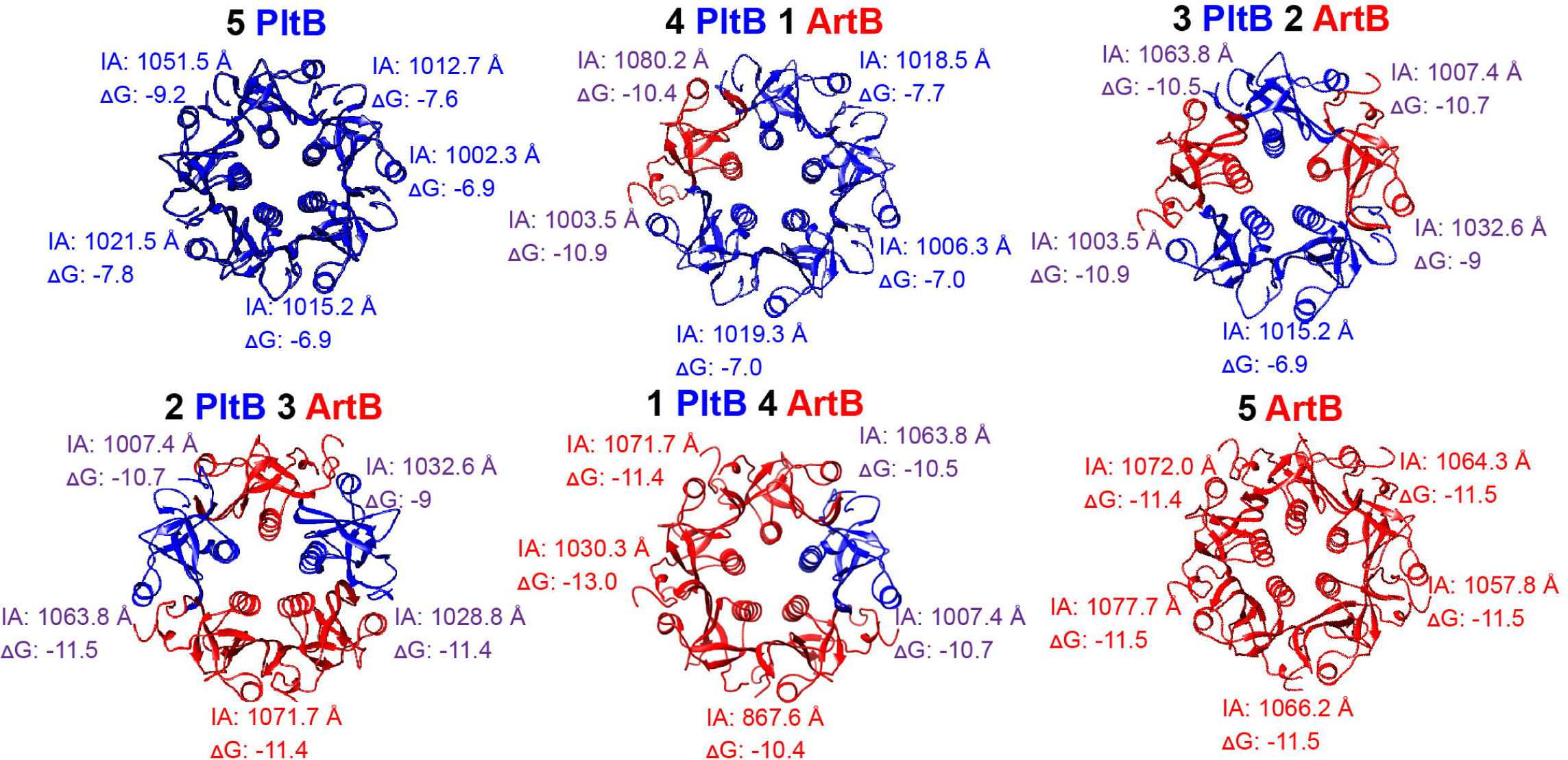
Modeling and interface analyses of binding subunits with different ratios of PltB and ArtB. Energies calculated between each binding subunit are color coded to represent the interaction: PltB-PltB are shown in blue, ArtB-ArtB in red, and PltB-ArtB in purple. IA: interface area in Angstroms (Å); ΔG solvation free energy gain upon formation of the interface (kcal/M). Negative ΔG values correspond to hydrophobic interfaces, and therefore a positive protein affinity.

**Figure 3.**
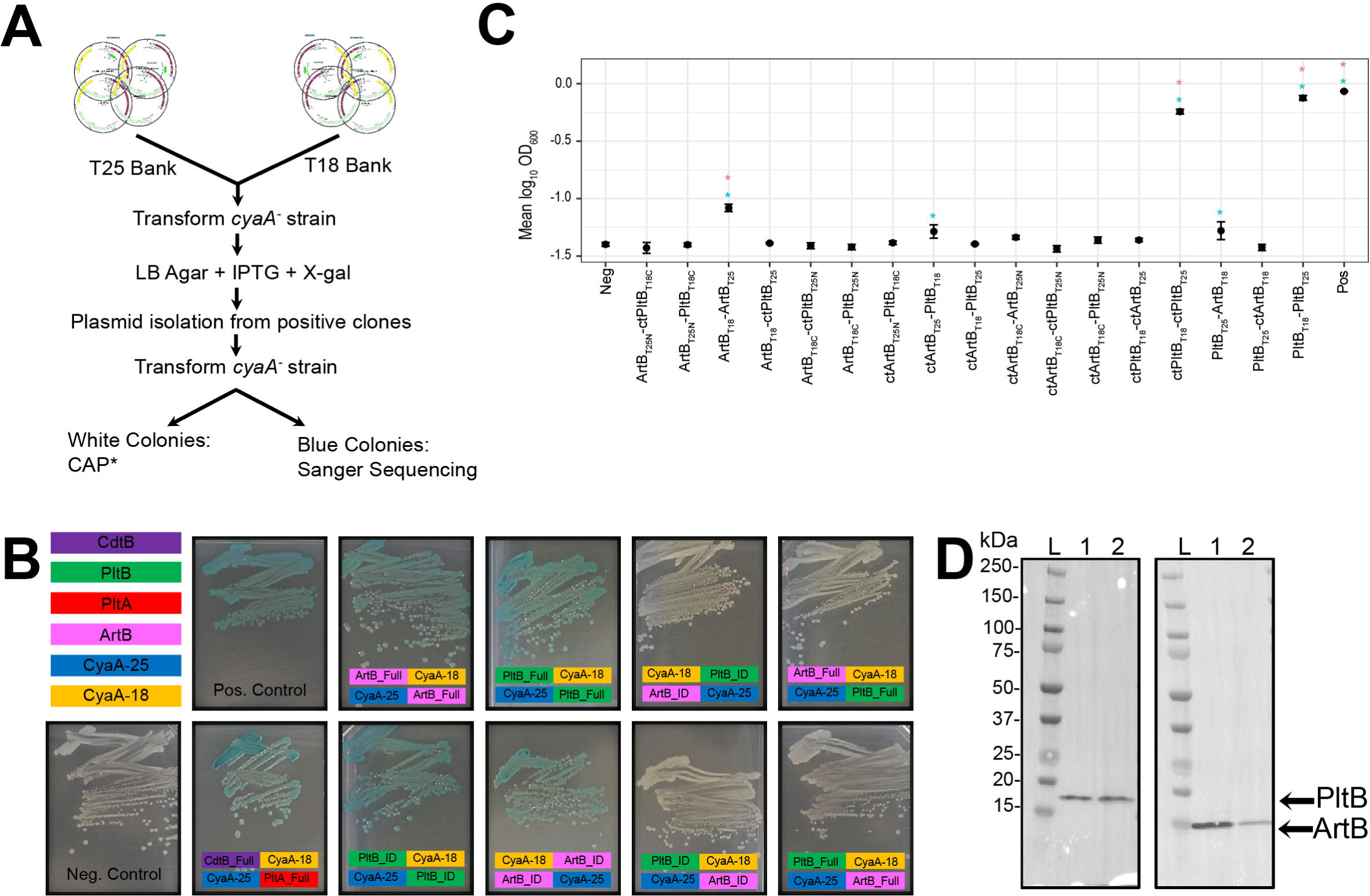
Analysis of PltB-ArtB heteropentamer formation. (**A**) Schematic of the method used to screen two-hybrid system clones. In the example shown, regions expressing different domains of PltB were fused to adenylate cyclase domain T18 or T25. All T18 or T25 clones were pooled into “banks” that were co-transformed into the cAMP-null (*cyaA*^-^) *E. coli* BTH101 strain. Screening was performed by growing the transformants on LB agar supplemented with IPTG and X-gal. Plasmid DNA was isolated from clones with positive interactions and was re-transformed into *E. coli* BTH101 to exclude the possibility of CAP* mutants. Confirmed interactions were submitted for Sanger sequencing to determine the identity of the interacting domains (**B**) *E. coli* BTH101 two-hybrid system strains grown on LB agar supplemented with IPTG and X-gal. Clones that showed positive interactions appear blue. Colored boxes show the fusion of either the interacting domain (_ID) or the full-length protein (_Full) for PltB, PltA, CdtB, and ArtB to either the CyaA-25 (shown in blue) or CyaA-18 subunits (shown in yellow) of the two-hybrid system. (**C**) Detection of interactionsof ArtB and PltB in *S.* Javiana *cyaA*^*-*^ strains harboring two-hybrid system plasmids. *S.* Javianacells were grown for 24 h in M63 minimal medium with maltose as the sole carbon source, and absorbance was measured at 600 nm. Error bars indicate the standard deviation of the mean of three independent experiments. Asterisks denote significant differences (p < 0.05) from the negative control before (blue) and after (pink) Dunnett’s test for multiple comparisons adjustment; the PltB_T25_ + ctArtB_T18_ interaction was marginally significant after multiple comparisons corrections (P = 0.0973). (**D**) Detection of PltB-3xFLAG and ArtB-c-Myc using TAP-tagging. Holotoxins were pulled down from *E. coli* BTH101 strains expressing CdtB-His, PltA-Strep, PltB-3xFLAG, and ArtB-c-Myc using anti-His antibodies (lane 1) followed by purification of PltB-3xFLAG-containing holotoxins by pulling down with anti-FLAG magnetic beads (lane 2). Proteins were visualized with antibody staining to detect PltB-3xFLAG (left panel) and ArtB-c-Myc (right panel). “L”represents the Western C (BioRad) protein ladder. The results of one representative experiment are shown; the assay was performed in two independent experiments.

### Construction of cyaA-S. Javiana

Deletion of *cyaA* was performed to enable screening for maltose utilization resulting from an interaction of T18- and T25-fused proteins. Whole genome sequence analysis identified a single full-length *cyaA* in the *S.* Javiana strain used in this study (FSL S5-0395). Construction of a *cyaA*^-^ *S.* Javiana strain was performed using the λ-Red recombinase system as described previously (Miller et al., 2018). Plasmids and primers used to generate the *cyaA*^*-*^strain are listed in Tables S2 and S3, respectively. The in-frame deletion was confirmed by Sanger sequencing. The mutant strain was also phenotypically confirmed by a lack of growth in M63 medium containing maltose as the sole carbon source; growth was restored upon addition of 0.05 mM cAMP.

### Expression of the two-hybrid system constructs in M63 minimal medium

Plasmids containing ArtB and PltB constructs were transformed into the *S.* Javiana *cyaA*^*-*^ strain via electroporation. Transformed cells were selected on LB agar plates supplemented with ampicillin and kanamycin. Overnight cultures (12 – 14 h) were grown shaking at 30°C in LB broth with ampicillin and kanamycin. Subsequently, M63 broth containing ampicillin and kanamycin, and 0.02 mM cAMP was inoculated with an overnight culture (diluted 1:100), followed by incubation with shaking at 30°C for 24 h. The addition of 0.02 mM cAMP (a concentration that does not support the growth of negative controls; Fig. 3C) was essential for the basal expression of the fusion proteins. Growth was assessed as optical density at 600 nm (OD_600_), which was measured after 24 h with a BioTek synergy plate reader (BioTek Instruments, Inc., Winooski, VT). Two-hybrid system interactions were also performed by growing *E. coli* BTH 101 cells harboring two-hybrid system plasmids in M63 broth containing ampicillin and kanamycin without exogenous cAMP. Growth was assessed using the same conditions as described for *S.* Javiana. Each growth assay was performed as three independent experiments for both *E. coli* and *S.* Javiana strains.

### Molecular modeling of different forms of the TT binding subunit

*In silico* modeling of TT subunits was done using the Phyre2 server (Kelley et al., 2015). Construction of the TT with different ratios of PltB and ArtB was performed using the MatchMaker tool in Chimera software (Pettersen et al., 2004). The stability and biological relevance of homo- and heteropentamer formation was calculated as free energies from individual subunit-subunit interactions within the pentamer, using the PDBe Protein Interfaces, Surfaces and Assemblies (PDBe PISA) server (Krissinel and Henrick, 2007).

### Tandem Affinity Purification (TAP)

Beveled flasks containing 250 mL of LB with ampicillin and kanamycin were inoculated with overnight cultures (12 – 14 h; 1:500 dilution) of *E. coli* BTH101 harboring plasmids pAG37 and pAG43, which express PltB-3xFlag ArtB-c-Myc, and CdtB-His PltA-Strep, respectively. Cells were grown shaking at 200 rpm for 4.5 h at 37°C, followed by induction with IPTG and cAMP (both 1 mM final concentration) for an additional 1.5 h. Cells were collected by centrifugation and stored at −80°C. Cells were thawed on ice, re-suspended in 2 ml of NTA buffer (50 mM NaH_2_PO_4_, pH 8.0, 0.3M NaCl) containing 2 mg lysozyme, and were incubated at 37°C for 30 min. Cell lysis was achieved with three rounds of freeze-thaw cycles (submersion in liquid nitrogen and incubation at 37°C) followed by sonication using a Branson Sonifier 250 sonicator (80% duty cycle, 7 output control) for 30 sec on ice (performed twice). Cell debris was removed by centrifugation at 15,000 x g for 10 min. For CdtB-His purification, 50 µl of Dynabeads (Thermo Fisher Scientific; Waltham, MA) were washed twice with 500 µl of cold NTA buffer in preparation for TAP. Cell lysates were added to beads, followed by incubation overnight (12 - 14 h) at 4°C in a tube rotator. Beads were collected on a magnetic stand and were washed three times at 4°C with 500 µl of cold NTA buffer with incubation periods of 10 min. Proteins were eluted using 300 µl of NTA buffer containing 250 mM imidazole. Eluted proteins were then dialyzed against TBS buffer (50 mM Tris-Cl, 150 mM NaCl, pH 7.5) containing 10% glycerol, and were divided into two fractions. PltB-3xFLAG was pulled down using anti-FLAG magnetic beads (Sigma-Aldrich) according to the manufacturer’s instructions. Proteins were eluted from beads by boiling in the presence of 100 µl of 1X SDS-loading dye. PltB-3xFLAG and ArtB-c-Myc were detected in sub-fractions using western blot analyses performed with rabbit anti-FLAG (Sigma-Aldrich) and rabbit anti-c-Myc (Sigma-Aldrich) antibodies.

### Western blot detection of proteins

Protein samples were resolved on 4-20% Mini-PROTEAN TGX Precast Protein SDS-PAGE gels (BioRad Laboratories; Hercules, CA) and blotted on PVDF membranes using the Trans-Blot Turbo transfer system (BioRad). Membranes were incubated in TBS containing 0.1% tween 20 (TTBS) and 5% blocking reagent (BioRad) with gentle shaking at room temperature for 30 min. Primary antibodies were added using the manufacturer’s recommended dilution in TTBS with 0.5% blocking reagent, followed by incubation with gentle shaking at room temperature for 12 – 14 h. Membranes were washed three times with TTBS, followed by incubation (2 h at room temperature with gentle shaking) with secondary antibodies added at manufacturer’s recommended dilutions in TTBS with 0.5% blocking reagent. Membranes were washed twice in TTBS and once in TBS, before detection. Horse radish peroxidase-conjugated secondary antibodies were detected using Clarity Western ECL substrate (BioRad). Blots were visualized using the BioRad ChemiDoc MP imaging system.

### Binding subunit exchange assay

Protein expression and cell lysis of *E. coli* BTH101 strains (FSL G4-0035 to G4-0038) were performed as described above for TAP with cell pellets resuspended in PBS containing 10% glycerol. Final total protein concentration was determined spectrophotometrically (Nanodrop 2000c) (Layne, 1957;Stoscheck, 1990). Total protein of the PltB-3xFLAG-containing lysate was added in varying concentrations to 100 µg of total lysate containing CdtB-His, PltA-FLAG, and ArtB-c-Myc for a final reaction volume of 50 µl of PBS with 10% glycerol. The reaction was incubated at 37°C for 30 min, followed by 2 h incubation at 4°C to ensure equilibrium. A 200 µl aliquot of NTA buffer was added and protein complexes were purified using Ni-NTA beads, as described above. The assay was also performed with lysates containing CdtB-His, PltA-FLAG, and PltB-3xFLAG as target and ArtB-c-Myc as the competitor. All proteins were detected with western blotting using antibodies that recognize the corresponding protein tag (i.e., anti-FLAG, anti-His, and anti-c-Myc); three independent experiments were performed.

### Intoxication of HIEC-6 cells with crude lysates of toxin components expressed in *E. coli* BTH101 cells

Fresh OptiMem (containing 10 ng/ml r-EGF and 10% FBS) medium supplemented with 10 µg/ml gentamicin was added to HIEC-6 cells grown to confluency on 12 mm coverslips (Thermo Fisher Scientific) in 24-well plates (Corning Inc.; Corning, NY). Lysates containing various combinations of toxin subunits were then added to HIEC-6 cells at a final concentration of 400 µg total protein per ml, and the HIEC-6 cells were subsequently incubated at 37°C with 4.5% CO_2_. Immunofluorescence (IF) detection of DDR foci and cell cycle analyses were performed at 24 ±2 h after inoculation. Three independent experiments were performed for both cell cycle and IF detection experiments.

### IF detection of DDR proteins γH2AX and 53BP1

IF staining for γH2AX and 53BP1 foci was performed as described previously (Miller and Wiedmann, 2016b;Miller et al., 2018). The following antibodies were used: mouse anti-γH2AX (EMD Millipore, Billerica, MA), rabbit anti-53BP1 (Novus Biologicals, Littleton, CO), donkey anti-rabbit conjugated to Alexa 555 (diluted 1:500), and donkey anti-mouse conjugated to Alexa 647 (1:200; all Thermo Fisher Scientific). Nuclei were stained with 4’,6-diamidino-2-phenylindole (DAPI; Thermo Fisher Scientific) for 5 min at room temperature. Microscopic observation was performed using a Zeiss 710 confocal microscope. FIJI software was used for image processing (Schindelin et al., 2012). Cells (at least 50 were scored per treatment) were considered positive if their nuclei had at least four 53BP1 foci, and also contained γH2AX foci.

### Cell cycle arrest

Staining with propidium iodide for cell cycle analysis determination was performed as described previously (Miller and Wiedmann, 2016b;Miller et al., 2018). DNA content (for cell cycle analysis) was assessed using the FACSARIA flow cytometer (BD). Gating was performed to exclude multiplets as described previously (Wersto et al., 2001;Miller et al., 2018).

### Phylogenetic analyses

Sequences from a convenience sample of 40 *S.* Javiana isolates, representing 28 unique SNP clusters were downloaded from the NCBI Pathogen Detection browser (https://www.ncbi.nlm.nih.gov/pathogens/isolates/#/search/; see Table S7 for details). *S.* Mississippi isolate SRR1960042 was included as an outgroup for phylogenetic analyses (see Table S7). Illumina adapters from sequence reads were trimmed, and low-quality bases were removed using Trimmomatic 0.33 with default settings (Bolger BM, 2014). Determination of the quality of trimmed reads was done using FastQC v0.11.7 (Andrews, 2010). *De novo* assembly of genomes was done using SPAdes 3.6.0 (Bankevich A, 2012). To assess the qualities of draft genomes, QUAST 3.2 (Gurevich A, 2013) was used, followed by Bbmap 35.49 (Busnell, 2015) and SAMtools 1.3.1 (Li H, 2009) to calculate the average coverage. Serotypes of draft genomes were confirmed using SISTR (Yoshida CE, 2016). kSNP3 was used to identify core SNPs in all 40 *S.* Javiana genomes and strain FSL S5-0395; a *k*-mer size of 19 was used, as determined using kSNP3’s Kchooser function (Gardner SN, 2015). A phylogenetic tree was constructed with RaXML (Stamatakis, 2014) using a general time-reversible model with gamma-distributed sites constructed from 1,000 bootstrap repetitions. FigTree v. 1.4.4 was used for editing RaXML output (Young et al., 2000).

### BLAST detection of TT genes and artAB in S. Javiana and amino acid alignment

The presence of *artA, artB, pltA, cdtB*, and *pltB* in all 40 *S.* Javiana isolates was determined using nucleotide BLAST (blastn) version 2.3.0 with a maximum e-value of 1e-20, a gap opening penalty of 3, and a gap extending penalty of 1, to query *artA, artB, pltA, cdtB*, and *pltB* sequences from *S.* Javiana strain CFSAN001992 (Allard et al., 2013) against the isolates’ assembled draft genomes (Camacho C, 2009). Geneious software (Auckland, New Zealand) was used to perform nucleotide and amino acid sequence alignments of the genes extracted from the 40 *S.* Javiana isolates.

### qPCR quantification of artB and pltB differential expression

Overnight cultures (16 – 18 h) of FSL S5-0395 grown in LB broth were sub-cultured 1:1000 into LB or N-salts minimal medium (either pH 7 or pH 5.8), and sub-cultured samples were grown at 37°C shaking at 200 rpm until cells reached mid-exponential phase (3 h for LB, 5 h for N-salts minimal medium). RNA was stabilized with RNA protect (Qiagen), and was collected using the RNEasy kit (Qiagen). DNA was depleted with Ambion Dnase I (Life Technologies), which was confirmed with qPCR; a cycle threshold of >34 for *rpoB* in Dnase-treated RNA samples was used as a threshold for successful DNA depletion. cDNA libraries were prepared with the Superscript Reverse Transcription kit (Thermofisher), according to manufacturer’s instructions. qPCR was performed with SYBR Green 2X Master Mix (Applied Biosystems) in a reaction containing 0.4 µM of each primer (see Table S4) and 1 µL of cDNA (∼approximately 15 ng of cDNA) as template. Fold expression was calculated by raising the ΔΔCt to the power of the efficiency calculated for each primer pair (Schmittgen and Livak, 2008). Results are the average of three independent experiments, each performed in technical duplicate.

### Statistical analyses and data availability

Statistical differences were assessed using R studio version 3.4.2. using packages lme4 (Bates et al.) 1.1-14, emmeans version 1.3.3 (2016), lmerTest version 2.0-33 (Kuznetsova et al., 2015), and multicomp version 1.4-8 (Hothorn et al., 2008) for Dunnett’s test for multiple comparisons adjustment. Scripts and data sets are available online at https://github.com/ram524/2019_ArtB.

## Results

### artB is highly conserved among S. Javiana isolates

As ArtB had previously been suggested to form a holotoxin with CdtB and PltA (Gao et al., 2017), we first determined the presence and conservation of ArtB and TT genes in *S.* Javiana (Fig. 1A), using BLAST searches for a sample of 40 isolates (Timme et al., 2019).All 40 *S.* Javiana isolates encoded full-length *pltA, pltB*, and *cdtB. artB* was detected in 38 of 40 (95%) *S.* Javiana isolates, with five of these 38 isolates (13.2%) containing a 46-nucleotide deletion in *artB* (shown in blue in Fig. 1B), resulting in a premature stop codon (Fig. 1C). Among the 33 isolates encoding a full-length *artB*, all had a 100% nucleotide identity over the full-length sequence of this gene (426 nt). In *S.* Javiana, ArtB, is encoded in an operon with *artA*, which includes a frame-shift mutation resulting in a premature stop codon (Fig.1A). The *artA* pseudogene was also detected among the 38 isolates that were confirmed to encode *artB*. As *artAB* was previously shown to be encoded on a prophage in *S.* Typhimurium DT104 (Saitoh et al., 2005), we also used (i) PHASTER (Arndt et al., 2016), and (ii) manual screening to check for prophage genes at or flanking the *artAB* locus for the closed *S.* Javiana genome CFSAN001992. Neither method detected prophage-encoded genes at the *artAB* locus, suggesting that *artAB* in *S.* Javiana is not encoded on a prophage.

### In silico modeling suggests heteropentamers of ArtB and PltB are energetically feasible

While the structure of the TT binding subunit with homopentamers of PltB (Song et al., 2013), and ArtB (Gao et al., 2017), have been resolved experimentally, the ability of the toxin to form heteropentamers was unknown. We first modeled the *S.* Javiana ArtB structure based on the *S*. Typhimurium DT104 ArtB structure and constructed models of the TT with different ratios of ArtB and PltB in the binding subunit. Interface areas and the ΔG solvation energy gain of the ArtB complex formation for *S.* Typhimurium DT104 ArtB and *S.* Javiana strain CFSAN001992 ArtB were highly similar, supporting the modeling strategy used. ArtB was shown to interact with more specificity and stronger hydrophobicity with itself (ArtB-ArtB ΔG average: −11.5) than with PltB (ArtB-PltB ΔG average: −10.5) (Fig. 2); however, the theoretical interaction of ArtB-PltB was predicted to be stronger than PltB-PltB (ΔG average:-7.5; Fig. 2).

### Two-hybrid system interactions suggest that ArtB and PltB interact in the native host S. Javiana, but not in E. coli BTH101

Next, we assessed the ArtB-PltB interaction using an adenylate-cyclase two-hybrid system in *E. coli* BTH101 in combination with a novel high-throughput screening method (Fig. 3A). We designed constructs to assess possible interaction domains for each toxin component (CdtB, PltA, PltB, and ArtB) including, (i) the full-length protein, (ii) polypeptide sequences without the signal sequence, and (iii) the oligomerization domain as predicted from the TT structure (Song et al., 2013). Using this technique, we confirmed PltB-PltB, ArtB-ArtB, and CdtB-PltA interactions (Fig. 3B), but a PltB-ArtB interaction was not detected (Fig. 3B and S1). As increasing levels of cAMP resulting from a positive interaction also activates the maltose catabolic pathway of the BACTH system, maltose utilization can also be used as a screening method as the use of maltose as the sole carbon source requires cAMP activation of the catabolite activator protein (Battesti and Bouveret, 2012). Therefore, we further confirmed the results of the two-hybrid system by growing a subset of the *E. coli* BTH101 cells in M63 minimal medium (without exogenous cAMP as this enabled growth of the negative control strain) and found that only one PltB-PltB clone showed significant growth, indicating an interaction; no ArtB-ArtB or PltB-ArtB interactions were detected.

To determine if an accessory protein, which may be absent in *E. coli* BTH101, was necessary for the interaction of ArtB and PltB, we constructed a *cyaA*^*-*^ *S.* Javiana mutant for testing interactions in *Salmonella* by transforming a sub-set of the PltB- and ArtB-containing constructs into *S.* Javiana *cyaA*^*-*^ strains, and then growing them in M63 minimal medium with maltose as the sole carbon source; growth in M63 minimal medium was used as an alternative screening method as *S.* Javiana is β-galactosidase null. In *S.* Javiana, there were significant (P < 0.05, uncorrected p-value) PltB-PltB, ArtB-ArtB, and ArtB-PltB interactions (see Fig. 3C and Table S6). Upon correcting for multiple comparisons, one of the ArtB-PltB interactions was significant at α = 0.1 (P = 0.0975; see Fig. 3C). Together, the results of the two-hybrid system suggest that ArtB-PltB interactions may occur in *S.* Javiana cells.

### PltB and ArtB proteins are co-purified and can form biologically active binding subunits of the TT

As previous studies have suggested that the TT binding subunit exists as a stable pentamer (Song et al., 2013), we reasoned that despite the weak ArtB-PltB interaction demonstrated by the two-hybrid system in *S.* Javiana, interactions in the assembled holotoxin might be more stable. Therefore, we used tandem affinity purification (TAP) to (i) determine if PltB and ArtB can interact with PltA and CdtB to form a complete holotoxin, and (ii) assess if holotoxins are formed with homo- or heteropentamers of PltB and ArtB. Purification of His-tagged CdtB followed by purification with 3xFLAG-tagged PltB (Fig. 3D) revealed that ArtB is co-purified with PltB in both pull-down steps, supporting that PltB and ArtB can form heteropentameric binding subunits.

To confirm the activity of both homo- and heteropentameric forms of the binding subunit, we co-incubated toxin subunits from the same lysates used for TAP with human intestinal epithelial cells (HIEC-6 cells). Co-incubation with the lysates containing all four toxin subunits (i.e. CdtB, PltA, PltB, and ArtB) resulted in approx. 83% of cells with an activated DDR (Fig. 4A and 4B). Similarly, HIEC-6 cells co-incubated with lysates containing CdtB and PltA with either PltB or ArtB, both resulted in DDR activation in approx. 78% of HIEC-6 cells. There was no difference (for all comparisons, P = 1) between lysates containing both PltB and ArtB subunits (which showed that ArtB was co-purified with PltB in TAP experiments; Fig. 3D), and those having only ArtB or only PltB homopentamers (Fig. 4A and 4B). Furthermore, cell cycle analyses of HIEC-6 cells co-incubated with lysates containing holotoxins with homopentamers of PltB or ArtB, or a mix of both PltB and ArtB binding subunits had a significantly higher (Mix: P = 0.005, PltB only: P = 0.037, and ArtB only: P = 0.006, respectively) proportion of cells accumulated in the G2/M phase, relative to control cells (Fig. 4C), a phenotype that is commonly associated with exposure to the TT (Haghjoo and Galán, 2004;Spanò et al., 2008).

**Figure 4.**
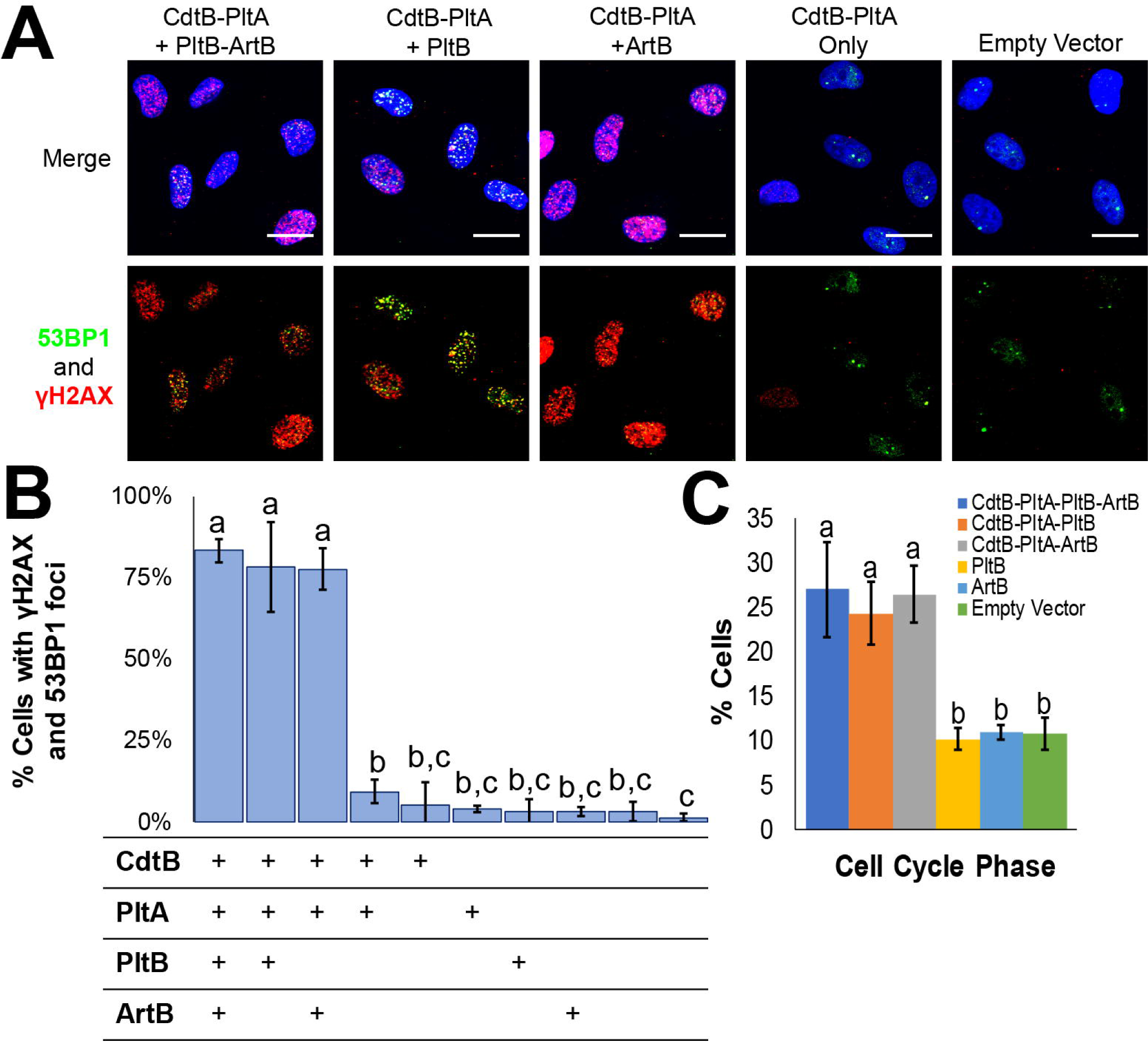
Holotoxins containing PltB-3xFLAG, ArtB-c-Myc, and a mix of PltB-3xFLAG and ArtB-c-Myc, activate the DDR in human intestinal epithelial cells (HIEC-6 cells). (**A**) IF staining of HIEC-6 cells co-incubated for 24 h with various total cell lysates containing CdtB-His, PltA-Strep, PltB-3xFLAG, and ArtB-c-Myc. DDR proteins are shown in green (53BP1) and red (γH2AX); nucleic acids are stained with DAPI (blue). Scale bar represents 20 µm. (**B**) Quantification of HIEC-6 cell nuclei having > four 53BP1-foci, as well as γ-H2AX foci; the table below the graph shows which subunits were present in the supernatant added to the HIEC-6 cells. Two negative controls were included and are represented by the final two bars in the graphic, which represent cell populations that were treated with supernatants from *E. coli* BTH101 cells with an empty vector (left) and untreated cell populations (right). (**C**) Proportion of cells in the G2/M cell cycle phase following co-incubation with different toxin subunit combinations for 24 hrs. Error bars represent standard deviations of the mean; bars that do not share letters are statistically different (P < 0.05). Results represent the average of three independent experiments.

### Both artB and pltB are co-expressed with cdtB at high levels in low Mg2+ medium

As *pltB* expression in *S.* Typhi (Fowler and Galan, 2018) was previously shown to occur when cells were cultured under Mg^2+^-limiting conditions, we compared the levels of RNA transcripts of *S.* Javiana cultured in LB broth (pH 7) and N-salts minimal medium containing 8 µM Mg^2+^ (pH 7). When *S.* Javiana was cultured in N-salts minimal medium (pH 7), RNA transcript levels of *pltB, cdtB*, and *artB* were on average 135-, 364-, and 44-fold higher, respectively, compared to levels in *S.* Javiana cells grown in LB broth. As production of the TT is hypothesized to occur when *Salmonella* is located within the *Salmonella*-containing vacuole (SCV) (Chang et al., 2016), we also grew *S.* Javiana in N-salts minimal medium acidified to pH 5.8, which has been shown previously to stimulate expression of SPI-2 genes (Deiwick et al., 1999). Under acidic conditions, expression of *cdtB* and *artB* increased marginally, to 382-fold and 86-fold (see Fig. 5B), while *pltB* transcript levels were lower when *S.* Javiana was cultured in N-salts minimal medium at pH 5.8 (approximately 2-fold lower compared to expression in N-salts minimal medium at pH 7). Finally, the ratio of ΔCt_*pltB*_: ΔCt_*artB*_ (representing the inverse relationship of the relative ratio of *pltB* transcripts to *artB* transcripts) was significantly lower for *S.* Javiana grown in N-minimal salts medium at pH 7 compared to pH 5.8 (p = 0.0047), suggesting that there are relatively higher levels of *pltB* transcripts at pH 7, than at pH 5.8. Together, these results suggest that *artB* and *pltB* are co-expressed with *cdtB* under low Mg^2+^-conditions, but *pltB* is expressed at relatively higher levels at neutral pH (pH 7), while *artB* is expressed at higher levels at pH 5.8.

**Figure 5.**
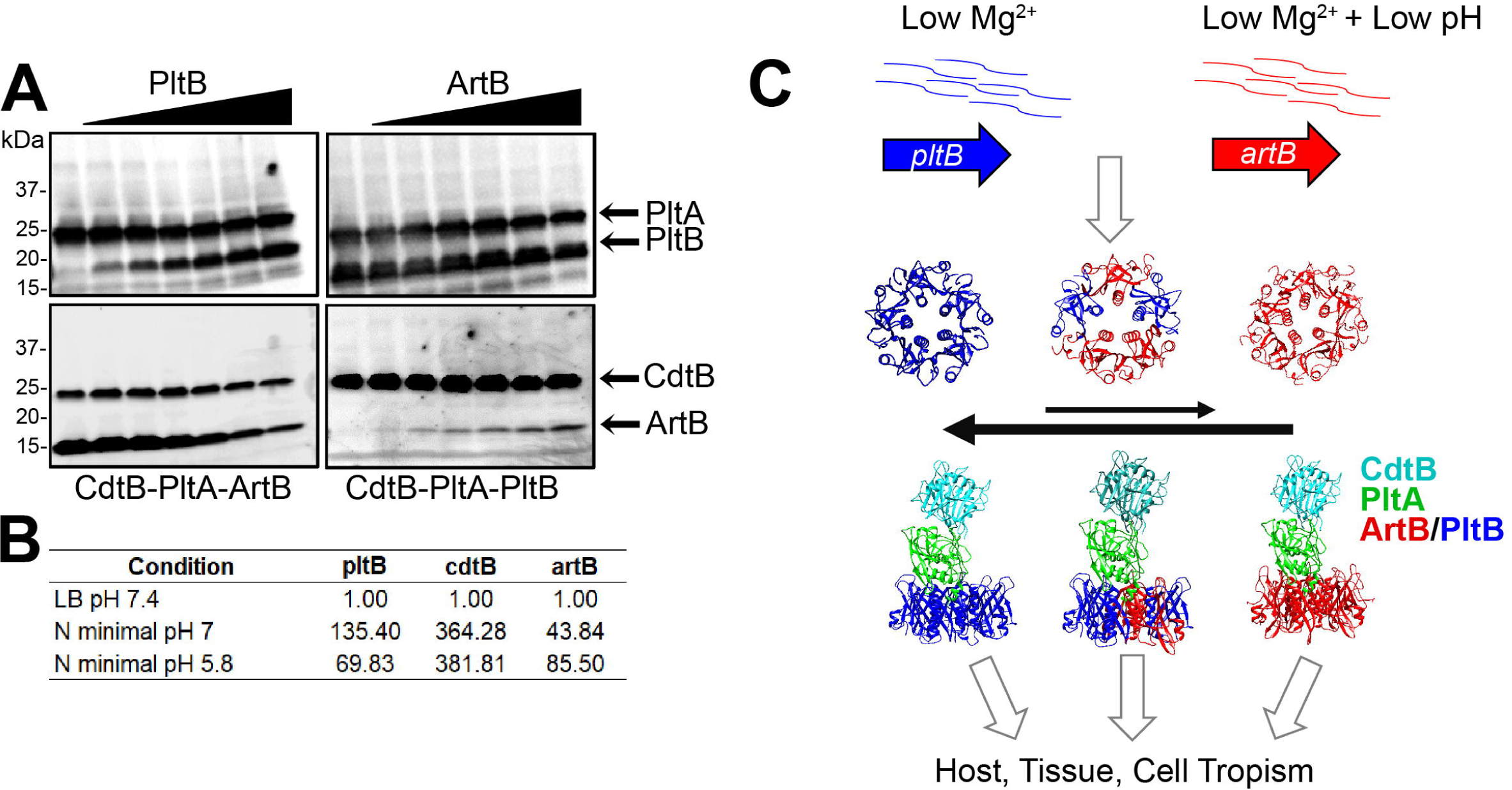
Competition and expression of PltB-ArtB heteropentamer formation *in vitro*. (**A**)Results of the binding subunit exchange assay. PltB-3xFLAG was added in varying concentrations to lysates containing CdtB-His, PltA-FLAG, ArtB-c-Myc (left panels), or ArtB-c-Myc was added to lysates containing CdtB-His, PltA-FLAG, PltB-3xFLAG (right panels). Holotoxins were purified by pulling down with anti-His to detect CdtB-His, and subunits were detected using tag-specific antibodies (PltA-FLAG, CdtB-His, and PltB-3xFLAG). One representative experiment is shown; the assay was performed in three independent experiments (**B**) Fold expression (2^-ΔΔCT^) of *pltB, cdtB*, and *artB* in *S.* Javiana cells grown for 5 h in either N salts minimal media at pH 7 or pH 5.8, normalized to expression of *S.* Javiana cells grown for 3 h in LB broth pH 7. Results are averaged from three independent experiments. (**C**) Proposed model for production of TT binding subunits. *artB* and *pltB* are co-expressed under low Mg^2+^ culturing conditions, but at neutral pH PltB outcompetes ArtB for inclusion in the final pentameric binding subunit, although homopentamers of ArtB and also heteropentamers of ArtB-PltB are also produced. The production of multiple variations of the binding subunit is predicted to expand the types of hosts, tissues, and cells that the toxin can bind to.

### PltB competes more efficiently for inclusion in the holotoxin than ArtB

Given that (i) *artB* and *pltB* are co-expressed, and (ii) ArtB and PltB are co-purified, we hypothesized that ArtB and PltB likely compete for inclusion in the holotoxin. Upon challenging the CdtB-PltA-ArtB complex with excess amounts of PltB followed by purification of the holotoxin, PltB efficiently replaced ArtB, thereby reducing the total amount of ArtB bound in the holotoxin (Fig. 5A). This result was not reciprocal, however, as ArtB was far less efficient at replacing PltB in a CdtB-PltA-PltB holotoxin (Fig. 5A). Together, this suggests that while ArtB can form a biologically-active binding subunit, PltB ultimately outcompetes for inclusion in the binding subunit.

## Discussion

Here, we show that the nontyphoidal serovar *S.* Javiana uses both ArtB and PltB to form homo-and heteropentameric binding subunits of the TT holotoxin. Interactions of ArtB and PltB are detected in *S.* Javiana cells, and ArtB and PltB are co-purified with CdtB and PltA (the active subunits of the toxin), indicating the formation of a heteropentameric holotoxin. Furthermore, *artB* and *pltB* are co-expressed along with *cdtB* under conditions which mimic the SCV (Deiwick et al., 1999). As a number of *Salmonella* serovars encode both *artB* and *pltB* (den Bakker et al., 2011;Rodriguez-Rivera et al., 2015), utilization of homo- and heteropentameric binding subunits suggests an evolutionary advantage for *Salmonella* serovars that encode both *artB* and *pltB*, as ArtB and PltB subunits have been shown to preferentially bind to different cells and tissues (Song et al., 2013;Gao et al., 2017).

### *artB* is generally conserved among *S.* Javiana isolates

Here we show that the majority of *S.* Javiana isolates also encode the *artAB* operon (95% of the isolates examined here). Interestingly, at least 25 other serovars have also been shown to encode *artAB* (Rodriguez-Rivera et al., 2015;Tamamura et al., 2017). In this study, some of the *S.* Javiana isolates harbored *artB* with a 46 bp internal deletion, which could reflect either acquisition of a mutated *artB* or slipped-strand mispairing that occurred during DNA replication, which has been reported previously for promoting phenotypic diversity in bacterial pathogens such as *B. pertussis* (Decker et al., 2012). Regardless, a high proportion of *S.* Javiana isolates encode a full-length *artB*, implicating that *artB* plays a role in the serovar’s virulence.

### ArtB and PltB are co-expressed, but PltB outcompetes for inclusion in the binding subunit

We previously showed that active TT was not produced by *S.* Javiana grown in LB broth (Miller and Wiedmann, 2016b). Here, we confirmed that expression of *pltB* is relatively low in LB broth, but expression is significantly induced (>100-fold) when *S.* Javiana cells are grown under Mg^2+^-limiting conditions, which have been shown to induce SPI-2 gene expression in *S.* Typhimurium (Deiwick et al., 1999;Fass and Groisman, 2009), and expression of *pltB* in *S.* Typhi (Fowler and Galan, 2018). Importantly, we also established that *artB* is co-expressed with *cdtB* and *pltB* in *S.* Javiana cells cultured under Mg^2+^-limiting conditions, despite *artB* being located nearly 500 kb upstream of the *cdtB*-islet (Fig. 1A).

Given that (i) *artB* and *pltB* are co-expressed, (ii) ArtB and PltB are structurally similar (Gao et al., 2017), and (iii) *in silico* analyses also suggested a favorable interaction between ArtB and PltB (Fig. 2), we hypothesized that ArtB and PltB might compete for inclusion in the binding subunit. While there was no evidence of an interaction between ArtB and PltB when the two-hybrid system was expressed in *E. coli* BTH101 cells, there was weak evidence to support an interaction in the native host *S.* Javiana (Fig. 3C). We were also able to detect ArtB in purified holotoxins using TAP to pull down CdtB, and then PltB, suggesting that ArtB and PltB interact. Furthermore, the ability of PltB to efficiently replace ArtB in a CdtB-PltA-ArtB holotoxin in a competition assay suggests that PltB has evolved as the preferred subunit of the TT, but some holotoxins likely contain a mixture of PltB and ArtB. Combined with evidence that suggests upregulation of *pltB* and *artB* under low Mg^2+^ conditions (even though the PltB-ArtB ratios seem to be modulated by pH), this evidence suggests that ArtB and PltB are co-expressed and hence may be incorporated into the holotoxin as either homo- or heteropentamers of ArtB and PltB subunits.

Although our data, in *S.* Javiana and *in vitro*, support a model in which ArtB and PltB compete for incorporation and formation of heteropentamers in the holotoxin, the inability to detect an ArtB-PltB interaction in *E. coli* BTH101 cells suggests that additional factors may contribute to the assembly of the binding subunit. Given that TT genes and *artB* are expressed when *Salmonella* cells are grown in low Mg^2+^ media, the requirement of a *Salmonella*-specific accessory protein such as a chaperone that is not expressed under the conditions used for the two-hybrid system in *E. coli*, or some other post-translational modification could explain why an interaction was observed in *S.* Javiana cells grown in M63 broth, but not in *E. coli* BTH101 cells.

While most AB5 toxin binding subunits exist as homopentamers of five identical monomers (i.e. shiga toxin, cholera toxin, subtilase toxin (Beddoe et al., 2010)), the pertussis toxin’s binding subunit, which is homologous to both the PltB subunit of the TT and ArtB, uses four distinct monomeric subunits (i.e., S2, S3, S4, and S5) to form the five-component heteropentameric binding subunit (Locht et al., 2011). *In vitro*, the pertussis toxin’s S2 subunit can be replaced by the S3 subunit (Raze et al., 2006). Our data suggest that ArtB and PltB likely compete for inclusion in the binding subunit in a manner similar to that of the pertussis toxin. Alternatively, or in addition, differential regulation might enable transcription of *artB* under select environmental conditions (e.g., under reduced pH and/or other conditions not tested here). Finally, while the theoretical modeling, which predicted stronger ArtB-ArtB hydrophobic interactions, suggested ArtB homopentamers would indeed be more energetically favorable, these models were done in absence of the PltA-CdtB subunits. It is possible that the interaction between PltA and the pentameric binding subunit is what ultimately determines the stability of the holotoxin, as PltA inserts into the barrel of the binding pentamer in the holotoxin (Song et al., 2013). For example, PltB homopentameric binding subunits might have a stronger interaction with PltA than with homo- or heteropentamers containing ArtB.

### Why express two binding subunits?

Why would NTS serovars encode multiple binding subunits? One possibility is that the ability to produce toxins with PltB or ArtB homopentamers or PltB-ArtB heteropentamers would expand the variety of cells, tissues, or even hosts that are susceptible to this toxin. While PltB has been reported to preferentially bind Neu5Ac-terminated glycans, which are abundantly present in human cells due to a mutation in the converting enzyme CMAH that is required to produce Neu5Gc-terminated glycans (Deng et al., 2014), ArtB preferentially binds Neu5Gc-terminated glycans (Gao et al., 2017). Neu5Ac (targeted by PltB) predominates in chicken, turkey, and fish (Samraj et al., 2015), while Neu5Gc-terminated glycan levels (targeted by ArtB) are high in cows, pigs, sheep, and goats (Samraj et al., 2015). As NTS serovars, including *S.* Javiana, are able to infect a broad host range, utilization of ArtB and PltB binding subunits would effectively expand the range of hosts that would be susceptible to this toxin. Another possibility is that *artB* and *pltB* are differentially expressed, as supported by our data that the ratio of *pltB*:*artB* is higher when cells are grown at neutral pH under Mg^2+^-limiting conditions, but *artB* is transcribed at relatively higher rates compared to *pltB* when cells are grown in acidified medium (pH 5.8). To date, the transcriptional and translational regulation of *artB* and *pltB*, and their corresponding gene products, have not been extensively characterized. Our data suggest that differential transcriptional regulation may lead to the production of different ratios of ArtB and PltB, allowing for different configurations of the binding subunit under different environmental conditions. For example, higher ArtB levels under acidic conditions may allow for enhanced ArtB levels in the acidified SCV.

## Conclusion

Overall, using *S.* Javiana as an example, our data support that ArtB and PltB compete for inclusion in the pentameric binding subunit of the TT, producing both homo- and heteropentameric forms of the binding subunit. As CDTs have been shown to play an important role in the long-term carriage of the toxin-producing pathogen (Ge et al., 2005;Del Bel Belluz et al., 2016), our data support an additional adaptation that could explain the broad host range of *S.* Javiana and other TT-positive NTS serovars (Hoelzer et al., 2011). Moreover, using the TT as a model to study AB5 toxins, our results suggest that the acquisition of multiple toxin binding subunits can be used as an evolutionary strategy to expand the number of cell, tissue, and host types that can be affected by the toxin.

## Acknowledgements

The authors gratefully acknowledge Dr. Pete Chandrangsu for his guidance with the *in silico* modelling, and the staff at the Cornell Statistical Consulting Unit and the Biotechnology Resource Center Imaging Facility for their guidance with statistical analyses, and flow cytometry data collection, respectively.

## Author Contributions

AG, ASH, MW, and RAC designed the study. AG, ASH, ARC, and RAC performed experimental, bioinformatical, and/or statistical analyses. All authors wrote and assisted with he revision of this manuscript.

