## Supplemental Tables for "The Typhoid Toxin Produced by the Nontyphoidal *Salmonella* Serovar Javiana Can Utilize Multiple Binding Subunits, which Compete for Inclusion in the Holotoxin"

**Table S1: Strains used in this study**

| **Strain** | **Relevant genotype/markers** | **Source** |
| --- | --- | --- |
| BTH101 | F^-^, *cya-99, araD139, galE15, galK16, rpsL1 (Str^r^ ), hsdR2, mcrA1, mcrB1* | (Karimova et al., 2001) |
| NEB5α | *fhuA2* Δ*(argF-lacZ)U169 phoA glnV44* Φ*80* Δ*(lacZ)M15 gyrA96 recA1 relA1 endA1 thi-1 hsdR17* | New England Biolabs |
| FSL G4-0003 | BTH101 containing pKT25 and pUT18C | This study |
| FSL G4-0004 | BTH101 containing pKT25-zip and pUT18C-zip | This study |
| FSL G4-0005 | BTH101 containing pAG37 and pAG40 | This study |
| FSL G4-0006 | BTH101 containing pAG37 and pAG43 | This study |
| FSL G4-0007 | BTH101 containing pAG40 | This study |
| FSL G4-0008 | BTH101 containing pAG38 | This study |
| FSL G4-0009 | BTH101 containing pAG39 | This study |
| FSL G4-0010 | BTH101 containing pAG41 | This study |
| FSL G4-0011 | BTH101 containing pAG42 | This study |
| FSL G4-0014 | BTH101 containing pAG23 and pAG25 | This study |
| FSL G4-0015 | BTH101 containing pAG19 and pAG21 | This study |
| FSL G4-0016 | BTH101 containing pAG29 and pAG31 | This study |
| FSL G4-0017 | BTH101 containing pAG34 and pAG36 | This study |
| FSL G4-0018 | BTH101 containing pAG25 and pAG29 | This study |
| FSL G4-0019 | BTH101 containing pAG25 and pAG33 | This study |
| FSL G4-0020 | BTH101 containing pAG19 and pAG31 | This study |
| FSL G4-0021 | BTH101 containing pAG19 and pAG35 | This study |
| FSL G4-0022 | BTH101 containing pAG21 and pAG33 | This study |
| FSL G4-0023 | BTH101 containing pAG21 and pAG29 | This study |
| FSL G4-0024 | BTH101 containing pAG23 and pAG31 | This study |
| FSL G4-0025 | BTH101 containing pAG19 and pAG35 | This study |
| FSL G4-0026 | BTH101 containing pAG24 and pAG36 | This study |
| FSL G4-0027 | BTH101 containing pAG20 and pAG36 | This study |
| FSL G4-0028 | BTH101 containing pAG25 and pAG34 | This study |
| FSL G4-0029 | BTH101 containing pAG22 and pAG34 | This study |
| FSL G4-0030 | BTH101 containing pAG24 and pAG32 | This study |
| FSL G4-0031 | BTH101 containing pAG26 and pAG32 | This study |
| FSL G4-0032 | BTH101 containing pAG26 and pAG30 | This study |
| FSL G4-0033 | BTH101 containing pAG22 and pAG30 | This study |
| FSL G4-0034 | BTH101 containing pAG5 and pAG9 | This study |
| FSL G4-0035 | BTH101 containing pAG40 and pAG38 | This study |
| FSL G4-0036 | BTH101 containing pAG40 and pAG39 | This study |
| FSL S5-0395 | *S.* Javiana wild-type | (Miller and Wiedmann, 2016) |
| FSL H9-0054 | *S.* Javiana *cyaA^-^* | This study |

**Table S2: Vectors and constructs used in this study**

| **Plasmid** | **Relevant genotype/markers** | **Source** |
| --- | --- | --- |
| pKT25 | Kan^R^, C-terminal fusion of adenylate cyclase T25 domain | (Karimova et al., 1998;Karimova et al., 2001) |
| pKNT25 | Kan^R^, N-terminal fusion of adenylate cyclase T25 domain | (Karimova et al., 1998;Karimova et al., 2001) |
| pUT18C | Amp^R^, C-terminal fusion of adenylate cyclase T18 domain | (Karimova et al., 1998;Karimova et al., 2001) |
| pUT18 | Amp^R^, N-terminal fusion of adenylate cyclase T18 domain | (Karimova et al., 1998;Karimova et al., 2001) |
| pKT25-zip | Kan^R^, positive control for adenylate cyclase two-hybrid system | (Karimova et al., 1998;Karimova et al., 2001) |
| pUT18C-zip | Amp^R^, positive control for adenylate cyclase two-hybrid system | (Karimova et al., 1998;Karimova et al., 2001) |
| pAG1 | pKNT25 containing full-length *cdtB* fused to adenylate cyclase T25 domain, Kan^R^ | This study |
| pAG2 | pUT18 containing full-length *cdtB* fused to adenylate cyclase T18 domain, Amp^R^ | This study |
| pAG3 | pKT25 containing *cdtB* lacking signal peptide coding region fused to adenylate cyclase T25 domain, Kan^R^ | This study |
| pAG4 | pKNT25 containing *cdtB* lacking signal peptide coding region fused to adenylate cyclase T25 domain, Kan^R^ | This study |
| pAG5 | pUT18 containing *cdtB* lacking signal peptide coding region fused to adenylate cyclase T18 domain, Amp^R^ | This study |
| pAG6 | pUT18C containing *cdtB* lacking signal peptide coding region fused to adenylate cyclase T18 domain, Amp^R^ | This study |
| pAG7 | pKNT25 containing full-length *pltA* fused to adenylate cyclase T25 domain, Kan^R^ | This study |
| pAG8 | pUT18 containing full-length *pltA* fused to adenylate cyclase T18 domain, Amp^R^ | This study |
| pAG9 | pKT25 containing *pltA* lacking signal peptide coding region fused to adenylate cyclase T25 domain, Kan^R^ | This study |
| pAG10 | pKNT25 containing *pltA* lacking signal peptide coding region fused to adenylate cyclase T25 domain, Kan^R^ | This study |
| pAG11 | pUT18 containing *pltA* lacking signal peptide coding region fused to adenylate cyclase T18 domain, Amp^R^ | This study |
| pAG12 | pUT18C containing *pltA* lacking signal peptide coding region fused to adenylate cyclase T18 domain, Amp^R^ | This study |
| pAG13 | pKT25 containing c-terminal region of *pltA* fused to adenylate cyclase T25 domain, Kan^R^ | This study |
| pAG14 | pKNT25 containing c-terminal region of *pltA* fused to adenylate cyclase T25 domain, Kan^R^ | This study |
| pAG15 | pUT18 containing c-terminal region of *pltA* fused to adenylate cyclase T18 domain, Amp^R^ | This study |
| pAG16 | pUT18C containing c-terminal region of *pltA* fused to adenylate cyclase T18 domain, Amp^R^ | This study |
| pAG17 | pKNT25 containing full-length *pltB* fused to adenylate cyclase T25 domain, Kan^R^ | This study |
| pAG18 | pUT18 containing full-length *pltB* fused to adenylate cyclase T18 domain, Amp^R^ | This study |
| pAG19 | pKT25 containing *pltB* lacking signal peptide coding region fused to adenylate cyclase T25 domain, Kan^R^ | This study |
| pAG20 | pKNT25 containing *pltB* lacking signal peptide coding region fused to adenylate cyclase T25 domain, Kan^R^ | This study |
| pAG21 | pUT18 containing *pltB* lacking signal peptide coding region fused to adenylate cyclase T18 domain, Amp^R^ | This study |
| pAG22 | pUT18C containing *pltB* lacking signal peptide coding region fused to adenylate cyclase T18 domain, Amp^R^ | This study |
| pAG23 | pKT25 containing c-terminal region of *pltB* fused to adenylate cyclase T25 domain, Kan^R^ | This study |
| pAG24 | pKNT25 containing c-terminal region of *pltB* fused to adenylate cyclase T25 domain, Kan^R^ | This study |
| pAG25 | pUT18 containing c-terminal region of *pltB* fused to adenylate cyclase T18 domain, Amp^R^ | This study |
| pAG26 | pUT18C containing c-terminal region of *pltB* fused to adenylate cyclase T18 domain, Amp^R^ | This study |
| pAG27 | pKNT25 containing full-length *artB* fused to adenylate cyclase T25 domain, Kan^R^ | This study |
| pAG28 | pUT18 containing full-length *artB* fused to adenylate cyclase T18 domain, Amp^R^ | This study |
| pAG29 | pKT25 containing *artB* lacking signal peptide coding region fused to adenylate cyclase T25 domain, Kan^R^ | This study |
| pAG30 | pKNT25 containing *artB* lacking signal peptide coding region fused to adenylate cyclase T25 domain, Kan^R^ | This study |
| pAG31 | pUT18 containing *artB* lacking signal peptide coding region fused to adenylate cyclase T18 domain, Amp^R^ | This study |
| pAG32 | pUT18C containing *artB* lacking signal peptide coding region fused to adenylate cyclase T18 domain, Amp^R^ | This study |
| pAG33 | pKT25 containing c-terminal region of *artB* fused to adenylate cyclase T25 domain, Kan^R^ | This study |
| pAG34 | pKNT25 containing c-terminal region of *artB* fused to adenylate cyclase T25 domain, Kan^R^ | This study |
| pAG35 | pUT18 containing c-terminal region of *artB* fused to adenylate cyclase T18 domain, Amp^R^ | This study |
| pAG36 | pUT18C containing c-terminal region of *artB* fused to adenylate cyclase T18 domain, Amp^R^ | This study |
| pAG37 | pUT18 containing *pltB-3x FLAG artB-c-Myc* | This study |
| pAG38 | pUT18 containing *pltB-3x FLAG* | This study |
| pAG39 | pUT18 containing *artB-c-Myc* | This study |
| pAG40 | pKNT25 containing *cdtB-His pltA-FLAG* | This study |
| pAG41 | pKNT25 containing *cdtB-His* | This study |
| pAG42 | pKNT25 containing *pltA-FLAG* | This study |
| pAG43 | pKNT25 containing *cdtB-His pltA-Strep* | This study |
| pKD46 | λ-Red recombinase plasmid | (Datsenko and Wanner, 2000) |
| pKD4 | Template plasmid | (Datsenko and Wanner, 2000) |
| pCP20 | Flippase plasmid | (Datsenko and Wanner, 2000) |

**Table S3: Primers used in this study**

| **Primer name** | **Sequence (5’ to 3’)** | **Usage** |
| --- | --- | --- |
| AG67-21-cdtB-BACTH-R | CTTAGGTACCCGACAGCTTCGTGCCAAAAAGGCT | Clone full-length *cdtB* in BACTH vectors |
| AG67-41-rbs-cdtB-F | CGCtctagaaTAGcgggagagtagatatcatgaa | Clone full-length *cdtB* in BACTH vectors |
| AG67-20-cdtB-BACTH-F | TCGACTCTAGAGAATATCAGTGACTACAAAGTTATG | Clone *cdtB* without signal sequence in BACTH vectors |
| AG67-21-cdtB-BACTH-R | CTTAGGTACCCGACAGCTTCGTGCCAAAAAGGCT | Clone *cdtB* without signal sequence in BACTH vectors |
| AG67-13-PltA-BACTH-R | CTTAGGTACCCGTTTAGAAAGTATAAGTTCTATTACA | Clone full-length *pltA* in BACTH vectors |
| AG67-37-rbs-pltA-F | CGCtctagaaTAGaggaGGAAGAAATAATGAAAAAGTTAATA | Clone full-length *pltA* in BACTH vectors |
| AG67-12-pltA-BACTH-F | TCGACTCTAGAGGTAGATTTTGTGTATCGTGTGGA | Clone *pltA* without signal sequence in BACTH vectors |
| AG67-13-PltA-BACTH-R | CTTAGGTACCCGTTTAGAAAGTATAAGTTCTATTACA | Clone *pltA* without signal sequence in BACTH vectors |
| AG67-39-rbs-artB-F | CGCtctagaaTAGaggaGgtggattatgaaaaagaaattaaag | Clone *pltA* C-terminal region in BACTH vectors |
| AG67-13-PltA-BACTH-R | CTTAGGTACCCGTTTAGAAAGTATAAGTTCTATTACA | Clone *pltA* C-terminal region in BACTH vectors |
| AG67-36-stop-pltB-F | GCGtctagaataATGTATATAAATAAGTTTGTGCCT | Clone full-length *pltB* in BACTH vectors |
| AG67-15-pltB-BACTH-R | CTTAGGTACCCGCTTGGGTCCAAAGCATTGTGT | Clone full-length *pltB* in BACTH vectors |
| AG67-14-PltB-BACTH-F | TCGACTCTAGAGGAGTGGACAGGAGATAAAACGA | Clone *pltB* without signal sequence in BACTH vectors |
| AG67-15-pltB-BACTH-R | CTTAGGTACCCGCTTGGGTCCAAAGCATTGTGT | Clone *pltB* without signal sequence in BACTH vectors |
| AG67-68-pltB-Ct-F | TCGACTCTAGAGAGCATATGGGCACCCTCCT | Clone *pltB* C-terminal region in BACTH vectors |
| AG67-15-pltB-BACTH-R | CTTAGGTACCCGCTTGGGTCCAAAGCATTGTGT | Clone *pltB* C-terminal region in BACTH vectors |
| AG67-39-rbs-artB-F | CGCtctagaaTAGaggaGgtggattatgaaaaagaaattaaag | Clone full-length *artB* in BACTH vectors |
| AG67-19-ArtB-BACTH-R | CTTAGGTACCCGATTTGTCAACATAGGCCCCATA | Clone full-length *artB* in BACTH vectors |
| AG67-18-ArtB-BACTH-F | TCGACTCTAGAGGCTATGGCTGATTATGATACGTA | Clone *artB* without signal sequence in BACTH vectors |
| AG67-19-ArtB-BACTH-R | CTTAGGTACCCGATTTGTCAACATAGGCCCCATA | Clone *artB* without signal sequence in BACTH vectors |
| AG67-67-ArtB-Ct-F | TCGACTCTAGAGGGGAATCATAAACAGGGGTTTG | Clone *artB* C-terminal region in BACTH vectors |
| AG67-19-ArtB-BACTH-R | CTTAGGTACCCGATTTGTCAACATAGGCCCCATA | Clone *artB* C-terminal region in BACTH vectors |
| AG67-35-rbs-pltB-F | cgctctagaAtaGATTTTAAATGTACAGGAGAGTA | Construction of *pltB-3x FLAG* |
| AG67-48-pltB-3xFLAG-R1 | ATCTTTATAATCACCATCATGATCTTTATAATCTGGACGCTTGGG  TCCAAAGCATTGTG | Construction of *pltB-3x FLAG* |
| AG67-49-pltB-3xFLAG-R2 | ATCATCATCTTTATAATCAATATCATGATCTTTATAATCACCATCATGATCT | Construction of *pltB-3x FLAG* |
| AG67-50-pltB-3xFLAG-R3 | CGCGGTACCTTATTTATCATCATCATCTTTATAATCAATATCATGATCTT | Construction of *pltB-3x FLAG* |
| AG67-51-artB-cmyc-F | CGCGGTACCATAGAGGAGGTGGATTATGAAAAAGAAATTAAAG | Construction of *artB-c-Myc* |
| AG67-52-artB-cmyc-R1 | TAATCAGTTTCTGTTCTGGACGATTTGTCAACATAGGCCCCAT | Construction of *artB-c-Myc* |
| AG67-53-artB-cmyc-R2 | CGCGAATTCTTACAGATCTTCTTCGCTAATCAGTTTCTGTTCTGGACGA | Construction of *artB-c-Myc* |
| AG67-41-rbs-cdtB-F | CGCtctagaaTAGcgggagagtagatatcatgaa | Construction of *cdtB-His* |
| AG67-54-cdtB-His-R1 | GATGATGATGATGATGTGGACGACAGCTTCGTGCCAAAAAGG | Construction of *cdtB-His* |
| AG67-55-cdtB-His-R2 | TATGGTACCTTAATGATGATGATGATGATGATGATGATGATGTGGAC | Construction of *cdtB-His* |
| AG67-56-pltA-FLAG-F | CGCGGTACCATAGAGGAGGAAGAAATAATGAAAAAGTTAATA | Construction of *pltA-FLAG* |
| AG67-57-pltA-FLAG-R1 | TCTTTATAATCTGGACGTTTAGAAAGTATAAGTTCTATTACA | Construction of *pltA-FLAG* |
| AG67-58-pltA-FLAG-R2 | CGCGAGCTCTTATTTATCATCATCATCTTTATAATCTGGACGTTTAGA | Construction of *pltA-FLAG* |
| AG67-56-pltA-FLAG-F | CGCGGTACCATAGAGGAGGAAGAAATAATGAAAAAGTTAATA | Construction of *pltA-Strep* |
| AG67-87-pltA-Strep-R1 | CGAACTGCGGGTGGCTCCAAGAACCTTTAGAAAGTATAAGTTCTATTACA | Construction of *pltA-Strep* |
| AG67-88-pltA-Strep-R2 | cgcgagctcttaTTTTTCGAACTGCGGGTGGCTCCA | Construction of *pltA-Strep* |
| SH144-cyaA-pKD4-F | TATTGAGACTCTGAAACAGAGACTGGATGTGTAGGCTGGAGCTGCTTC | *cyaA* in-frame deletion |
| SH145-cyaA-pKD4-R | ATACTGCTGCAATAGCGGCGCGTCATGATCCATATGAATATCCTCCTTAG | *cyaA* in-frame deletion |
| SH126-dcyaA-F | TTTAAGAATTTACACGCAGCGAACGGTGCTAC | Sequence of *cyaA* mutant |
| SH127-dcyaA-R | ACGCATCTGCCACCTCAAACCTCCTCCTGTAAA | Sequence of *cyaA* mutant |
| RM79_cdtB_RTPCR_F | ACCTGGAATCTCGGAACCAC | qPCR detection of *cdtB* |
| RM80_cdtB_RTPCR_R | GGCGAGATGCGACAGTTGTC | qPCR detection of *cdtB* |
| RM83_pltB_RTPCR_F | GGTGTACCGCTACCGTTAGC | qPCR detection of *pltB* |
| RM84_pltB_RTPCR_R | TTAATGCCAGCGCTGAGTGG | qPCR detection of *pltB* |
| RM85_rpoB_RTPCR_F | TACGGGACGCATCCACGTAC | qPCR detection of *rpoB* |
| RM86_rpoB_RTPCR_R | GTGCGAACATGCAACGTCAG | qPCR detection of *rpoB* |
| RM298_artBqPCR_F | GAGTGAGGCTATTAGTGTCAATGCCAT | qPCR detection of *artB* |
| RM299_artBqPCR_R | TCCCTGGAAGAGAATGCACTTTTGA | qPCR detection of *artB* |

**Table S4: Plasmids pooled into banks**

| Banks | Plasmids | No. of plasmids |
| --- | --- | --- |
| PltB-T18 | pAG18, pAG21, pAG22, pAG25, pAG26 | 5 |
| PltB-T25 | pAG17, pAG19, pAG20, pAG23, pAG24 | 5 |
| PltA-T18 | pAG8, pAG11, pAG12, pAG15, pAG16 | 5 |
| PltA-T25 | pAG7, pAG9, pAG10, pAG13, pAG14 | 5 |
| CdtB-T18 | pAG2, pAG5, pAG6 | 3 |
| CdtB-T25 | pAG1, pAG3, pAG4 | 3 |
| ArtB-T18 | pAG28, pAG31, pAG32, pAG35, pAG36 | 5 |
| ArtB-T25 | pAG27, pAG29, pAG30, pAG33, pAG34 | 5 |

**Table S5: Bank combinations transformed into *E. coli* BTH101**

| **Combination of banks** | **No. of combinations** |
| --- | --- |
| PltB+ArtB |  |
| T18+T25 | 50 |
| T18+T25 |  |
| PltB+PltA |  |
| T18+T25 | 50 |
| T18+T25 |  |
| CdtB+PltA |  |
| T18+T25 | 30 |
| T18+T25 |  |
| ArtB+PltA |  |
| T18+T25 | 50 |
| T18+T25 |  |
| PltB+PltB |  |
| T18+T25 | 25 |
| ArtB+ArtB |  |
| T18+T25 | 25 |
| Total | 230 |

**Table S6: Estimates of linear model on log10 transformed absorbance (OD_600_) data of *S*. Javiana *cyaA^-^* mutants**

| Plasmid combinations^a^ | Estimate | Standard Error | t value | p value | p value adjust^b^ |
| --- | --- | --- | --- | --- | --- |
| ArtB_T25N_ + ctPltB_T18C_ | -0.030360 | 0.043426 | -0.699 | 0.48897 | 0.9995 |
| ArtB_T25N_ + PltB_T18C_ | -0.003575 | 0.043426 | -0.082 | 0.93485 | 1.0000 |
| ArtB_T18_ + ArtB_T25_ | 0.317006 | 0.043426 | 7.300 | 1.33e-08 | <0.001 |
| ArtB_T18_ + ctPltB_T25_ | 0.011286 | 0.043426 | 0.260 | 0.79643 | 1.0000 |
| ArtB_T18C_ + ctPltB_T25_ | -0.011878 | 0.043426 | -0.274 | 0.78602 | 1.0000 |
| ArtB_T18C_ + PltB_T25_ | -0.022905 | 0.043426 | -0.527 | 0.60112 | 1.0000 |
| ctArtB_T25_ + PltB_T18C_ | 0.014355 | 0.043426 | 0.331 | 0.74289 | 1.0000 |
| ctArtB_T25_ + PltB_T18_ | 0.112489 | 0.043426 | 2.590 | 0.01376 | 0.1353 |
| ctArtB_T18_ + PltB_T25_ | 0.004132 | 0.043426 | 0.095 | 0.92472 | 1.0000 |
| ctArtB_T18C_ + ArtB_NT25_ | 0.060729 | 0.043426 | 1.398 | 0.17054 | 0.8166 |
| ctArtB_T18C_ + ctPltB_NT25_ | -0.038878 | 0.043426 | -0.895 | 0.37659 | 0.9924 |
| ctArtB_T18C_ + PltB_NT25_ | 0.037090 | 0.043426 | 0.854 | 0.39870 | 0.9952 |
| ctPltB_T18_ + ctArtB_T25_ | 0.038253 | 0.043426 | 0.881 | 0.38423 | 0.9935 |
| ctPltB_T18_ + ctPltB_T25_ | 1.155483 | 0.043426 | 26.608 | < 2e-16 | <0.001 |
| PltB_T25_ + ArtB_T18_ | 0.119244 | 0.043426 | 2.746 | 0.00936 | 0.0973 |
| PltB_T25_ + ctArtB_T18_ | -0.027102 | 0.043426 | -0.624 | 0.53651 | 0.9999 |
| PltB_T18_ + PltB_T25_ | 1.274816 | 0.043426 | 29.356 | < 2e-16 | <0.001 |
| Pos | 1.333219 | 0.043426 | 30.701 | < 2e-16 | <0.001 |

^a^subunits are abbreviated as T25 and T18 for the CyaA fusion where “T25” indicates C-terminal fusion, “T25N” N-terminal fusion, “T18” N-terminal fusion and “T18C” C-terminal fusion to the subunit; ct refers to the C-terminal of the typhoid toxin protein

^b^p value after multiple comparison adjustment with Dunnett contrasts

| **Table S7. SRR run identity and metadata for 40 *S.* Javiana isolates and one *S.* Mississippi isolate used for phylogenetic and BLAST analyses** | | | | | |
| --- | --- | --- | --- | --- | --- |
| **Serovar^a^** | **Run #^b^** | **Source** | **Country^c^** | **SNP Cluster^d^** | **Reason for Inclusion** |
| Javiana | SRR7081896 | Human clinical (stool) | USA | Not listed | Clinical source |
| Javiana | SRR7063447 | Human clinical (blood) | USA | [PDS000026970.5](https://www.ncbi.nlm.nih.gov/Structure/tree/#!/tree/Salmonella/PDG000000002.1290/PDS000026970.59?accessions=PDT000309089.2) | Clinical source |
| Javiana | SRR7063441 | Human clinical (stool) | USA | [PDS000026970.5](https://www.ncbi.nlm.nih.gov/Structure/tree/#!/tree/Salmonella/PDG000000002.1290/PDS000026970.59?accessions=PDT000309083.2) | Clinical source |
| Javiana | SRR7063436 | Human clinical (stool) | USA | [PDS000026970.5](https://www.ncbi.nlm.nih.gov/Structure/tree/#!/tree/Salmonella/PDG000000002.1290/PDS000026970.59?accessions=PDT000309078.2) | Clinical source |
| Javiana | SRR5816109 | Animal (swine) | USA | [PDS000026915.1](https://www.ncbi.nlm.nih.gov/Structure/tree/#!/tree/Salmonella/PDG000000002.1290/PDS000026915.13?accessions=PDT000224375.2) | Unique source |
| Javiana | SRR6311987 | Animal (canine) | USA | [PDS000036032.1](https://www.ncbi.nlm.nih.gov/Structure/tree/#!/tree/Salmonella/PDG000000002.1290/PDS000036032.1?accessions=PDT000266003.2) | Unique source |
| Javiana | SRR5598907 | Environmental (frog legs) | USA | [PDS000027056.1](https://www.ncbi.nlm.nih.gov/Structure/tree/#!/tree/Salmonella/PDG000000002.1290/PDS000027056.1?accessions=PDT000212911.2) | Unique source |
| Javiana | SRR2585797 | Environmental (toad) | Mexico | [PDS000027026.3](https://www.ncbi.nlm.nih.gov/Structure/tree/#!/tree/Salmonella/PDG000000002.1290/PDS000027026.3?accessions=PDT000086660.2) | Unique source |
| Javiana | SRR2968972 | Environmental (creek water) | USA | [PDS000026970.59](https://www.ncbi.nlm.nih.gov/Structure/tree/#!/tree/Salmonella/PDG000000002.1290/PDS000026970.59?accessions=PDT000095591.2) | Unique source |
| Javiana | SRR4124058 | Human clinical (stool) | USA | Not listed | Clinical source |
| Javiana | SRR6480222 | Human clinical (stool) | USA | [PDS000002099.77](https://www.ncbi.nlm.nih.gov/Structure/tree/#!/tree/Salmonella/PDG000000002.1290/PDS000002099.77?accessions=PDT000278865.2) | Clinical source |
| Javiana | SRR6396486 | Human clinical (stool) | USA | [PDS000026970.59](https://www.ncbi.nlm.nih.gov/Structure/tree/#!/tree/Salmonella/PDG000000002.1290/PDS000026970.59?accessions=PDT000273569.2) | Clinical source |
| Javiana | SRR6321401 | Human clinical (stool) | USA | [PDS000002101.29](https://www.ncbi.nlm.nih.gov/Structure/tree/#!/tree/Salmonella/PDG000000002.1290/PDS000002101.29?accessions=PDT000266798.2) | Unique SNP cluster |
| Javiana | SRR5724832 | Human clinical (stool) | USA | [PDS000002099.77](https://www.ncbi.nlm.nih.gov/Structure/tree/#!/tree/Salmonella/PDG000000002.1290/PDS000002099.77?accessions=PDT000219749.2) | Clinical source |
| Javiana | SRR5676348 | Human clinical (stool) | USA | [PDS000026970.59](https://www.ncbi.nlm.nih.gov/Structure/tree/#!/tree/Salmonella/PDG000000002.1290/PDS000026970.59?accessions=PDT000218237.2) | Clinical source |
| Javiana | SRR5605812 | Human clinical (urine) | USA | [PDS000026970.59](https://www.ncbi.nlm.nih.gov/Structure/tree/#!/tree/Salmonella/PDG000000002.1290/PDS000026970.59?accessions=PDT000213038.2) | Clinical source |
| Javiana | SRR1509381 | Human clinical (stool) | USA | [PDS000031655.1](https://www.ncbi.nlm.nih.gov/Structure/tree/#!/tree/Salmonella/PDG000000002.1290/PDS000031655.1?accessions=PDT000039538.2) | Unique SNP cluster |
| Javiana | SRR1561167 | Food (lobster) | Indonesia | Not listed | Unique location |
| Javiana | SRR1962283 | Human clinical | UK | Not listed | Clinical source |
| Javiana | SRR1965602 | Human clinical | UK | [PDS000031042.1](https://www.ncbi.nlm.nih.gov/Structure/tree/#!/tree/Salmonella/PDG000000002.1290/PDS000031042.1?accessions=PDT000055275.2) | Unique SNP cluster |
| Javiana | SRR3092111 | Food (peppers) | Mexico | Not listed | Unique source |
| Javiana | SRR3137186 | Human clinical (stool) | USA | [PDS000026970.59](https://www.ncbi.nlm.nih.gov/Structure/tree/#!/tree/Salmonella/PDG000000002.1290/PDS000026970.59?accessions=PDT000104780.2) | Clinical source |
| Javiana | SRR5152112 | Human clinical (stool) | USA | [PDS000030196.1](https://www.ncbi.nlm.nih.gov/Structure/tree/#!/tree/Salmonella/PDG000000002.1290/PDS000030196.1?accessions=PDT000176547.2) | Unique SNP cluster |
| Javiana | SRR5160225 | Human clinical (stool) | USA | [PDS000027927.45](https://www.ncbi.nlm.nih.gov/Structure/tree/#!/tree/Salmonella/PDG000000002.1290/PDS000027927.45?accessions=PDT000176928.2) | Unique SNP cluster |
| Javiana | SRR5289505 | Food (candy) | Pakistan | [PDS000001767.5](https://www.ncbi.nlm.nih.gov/Structure/tree/#!/tree/Salmonella/PDG000000002.1290/PDS000001767.5?accessions=PDT000190154.2) | Unique location |
| Javiana | SRR5429767 | Human clinical (stool) | USA | [PDS000002100.30](https://www.ncbi.nlm.nih.gov/Structure/tree/#!/tree/Salmonella/PDG000000002.1290/PDS000002100.30?accessions=PDT000200587.2) | Unique SNP cluster |
| Javiana | SRR5508324 | Human clinical (stool) | USA | [PDS000002089.70](https://www.ncbi.nlm.nih.gov/Structure/tree/#!/tree/Salmonella/PDG000000002.1290/PDS000002089.70?accessions=PDT000207146.2) | Unique SNP cluster |
| Javiana | SRR5583207 | Human clinical | UK | [PDS000028979.2](https://www.ncbi.nlm.nih.gov/Structure/tree/#!/tree/Salmonella/PDG000000002.1290/PDS000028979.2?accessions=PDT000211787.2) | Unique SNP cluster |
| Javiana | SRR5725238 | Human clinical (stool) | USA | [PDS000026937.9](https://www.ncbi.nlm.nih.gov/Structure/tree/#!/tree/Salmonella/PDG000000002.1290/PDS000026937.9?accessions=PDT000219854.2) | Unique SNP cluster |
| Javiana | SRR7346572 | Human clinical (stool) | USA | [PDS000026970.58](https://www.ncbi.nlm.nih.gov/Structure/tree/#!/tree/Salmonella/PDG000000002.1289/PDS000026970.58?accessions=PDT000329926.1) | Unique SNP cluster |
| Javiana | SRR6791758 | Human clinical (stool) | USA | [PDS000027037.34](https://www.ncbi.nlm.nih.gov/Structure/tree/#!/tree/Salmonella/PDG000000002.1290/PDS000027037.34?accessions=PDT000290255.2) | Unique SNP cluster |
| Javiana | SRR5217754 | Human clinical (stool) | USA | [PDS000026970.59](https://www.ncbi.nlm.nih.gov/Structure/tree/#!/tree/Salmonella/PDG000000002.1290/PDS000026970.59?term=serovar%3A%22Javiana%22) | Clinical source |
| Javiana | SRR5217761 | Human clinical (stool) | USA | [PDS000026982.28](https://www.ncbi.nlm.nih.gov/Structure/tree/#!/tree/Salmonella/PDG000000002.1290/PDS000026982.28?term=serovar%3A%22Javiana%22) | Unique SNP cluster |
| Javiana | SRR6425474 | Human clinical (stool) | USA | [PDS000027004.27](https://www.ncbi.nlm.nih.gov/Structure/tree/#!/tree/Salmonella/PDG000000002.1290/PDS000027004.27?term=serovar%3A%22Javiana%22) | Unique SNP cluster |
| Javiana | SRR5209739 | Human clinical (blood) | USA | [PDS000026915.13](https://www.ncbi.nlm.nih.gov/Structure/tree/#!/tree/Salmonella/PDG000000002.1290/PDS000026915.13?term=serovar%3A%22Javiana%22) | Unique SNP cluster |
| Javiana | SRR5023225 | Human clinical (stool) | USA | [PDS000026926.16](https://www.ncbi.nlm.nih.gov/Structure/tree/#!/tree/Salmonella/PDG000000002.1290/PDS000026926.16?term=serovar%3A%22Javiana%22) | Unique SNP cluster |
| Javiana | SRR6760627 | Human clinical (stool) | USA | [PDS000026948.5](https://www.ncbi.nlm.nih.gov/Structure/tree/#!/tree/Salmonella/PDG000000002.1290/PDS000026948.5?term=serovar%3A%22Javiana%22) | Unique SNP cluster |
| Javiana | SRR2601212 | Human clinical (stool) | USA | [PDS000027935.2](https://www.ncbi.nlm.nih.gov/Structure/tree/#!/tree/Salmonella/PDG000000002.1290/PDS000027935.2?term=serovar%3A%22Javiana%22) | Unique SNP cluster |
| Javiana | SRR1754882 | Human clinical (stool) | USA | [PDS000031372.5](https://www.ncbi.nlm.nih.gov/Structure/tree/#!/tree/Salmonella/PDG000000002.1290/PDS000031372.5?term=serovar%3A%22Javiana%22) | Unique SNP cluster |
| Javiana | SRR8082595 | Food (kratom) | USA | [PDS000027723.3](https://www.ncbi.nlm.nih.gov/Structure/tree/#!/tree/Salmonella/PDG000000002.1290/PDS000027723.3?term=serovar%3A%22Javiana%22) | Unique source |
| Mississippi | SRR1960042 | Human clinical | UK | PDS000006090.1 | Outgroup |

^a^Serovar confirmed with SISTR program

^b^SRR run ID assigned by NCBI

^c^Country of origin

^d^SNP cluster assigned by NCBI’s Pathogen Browser website as of 03/21/2019
