## Supplemental Figure Caption for "The Typhoid Toxin Produced by the Nontyphoidal *Salmonella* Serovar Javiana Can Utilize Multiple Binding Subunits, which Compete for Inclusion in the Holotoxin"

**Supplementary Figure Legend**

**Figure S1.** Results of two-hybrid system interactions in *E. coli* BTH101 cells. Cells were transformed with plasmids encoding partial interacting domain (_ID) or full-length proteins (_Full) fused to T18 and T25 subunits of CyaA (Karimova et al., 2001). Blue pigmentation suggests an interaction, while white colonies are unable to confirm an interaction.
