## Supplementary figures and images for "The Typhoid Toxin Produced by the Nontyphoidal *Salmonella* Serovar Javiana Can Utilize Multiple Binding Subunits, which Compete for Inclusion in the Holotoxin"

### Supplemental Figure 1

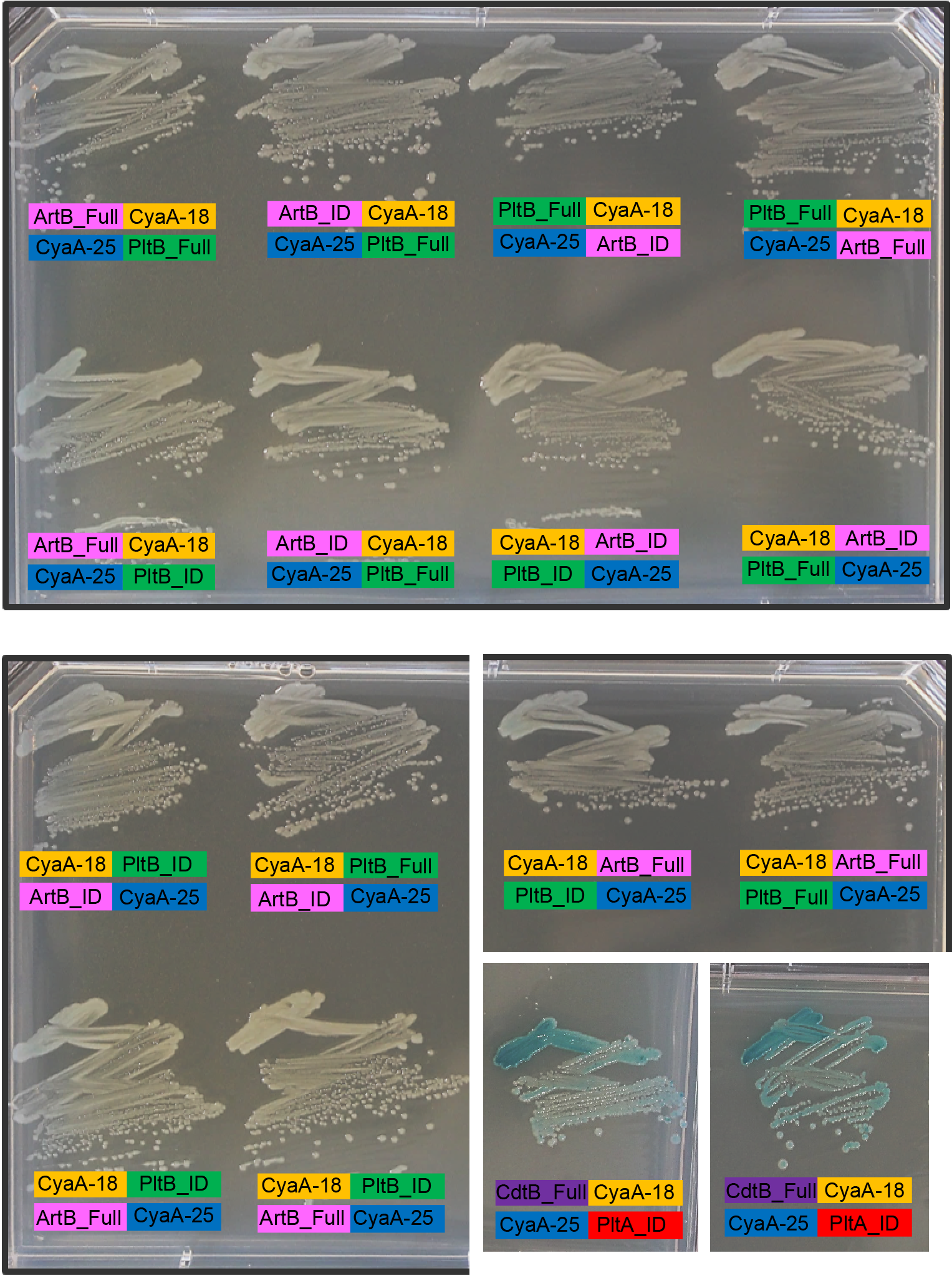
